# Regulation of lipid dysmetabolism and neuroinflammation progression linked with Alzheimer’s disease through modulation of Dgat2

**DOI:** 10.1101/2025.02.18.638929

**Authors:** Archana Yadav, Xiaosen Ouyang, Morgan Barkley, John C Watson, Kishore Madamanchi, Josh Kramer, Jianhua Zhang, Girish Melkani

**Author notes:** Corresponding Address: Department of Pathology, Division of Molecular and Cellular Pathology, School of Medicine, University of Alabama at Birmingham, AL 35294, USA.; (GCM).

## Abstract

Alzheimer’s disease (AD), an age-associated neurodegenerative disorder, is characterized by progressive cognitive decline, amyloid-β (Aβ) deposition, lipid dysregulation, and neuroinflammation. Although mutations in the amyloid precursor protein (APP) and accumulation of Aβ42 are established drivers of pathology, the mechanisms connecting amyloid toxicity with lipid metabolism and inflammatory responses remain poorly understood. Here, we employed complementary Drosophila and mouse models to dissect these relationships. Expression of App^NLG^ and Aβ42 in Drosophila resulted in locomotor deficits, disrupted sleep–circadian rhythms, memory impairments, lipid accumulation, synaptic loss, and neuroinflammatory signatures. Comparable lipid metabolic disturbances and inflammatory alterations were detected in the App^NLG-F^ knock-in mouse model, underscoring their conserved relevance to AD pathogenesis. We further identified diacylglycerol O-acyltransferase 2 (Dgat2), a key enzyme catalyzing the final step of triglyceride synthesis, as a critical modulator of AD-related phenotypes. Dgat2 expression was altered in both animal models and human AD tissues. Notably, panneuronal knockdown of Dgat2 in Drosophila attenuated lipid accumulation, restored synaptic integrity, and ameliorated locomotor and cognitive deficits, while also reducing neuroinflammation. Dgat2 suppression additionally improved sleep and circadian behavior, highlighting its pleiotropic protective effects. Together, these findings demonstrate a mechanistic link between amyloid pathology, lipid dysregulation, and neuroinflammatory processes. Targeting Dgat2 may therefore represent a novel therapeutic strategy to counteract AD-associated metabolic and neuronal dysfunction. The conservation of lipid homeostasis mechanisms across species underscores the translational potential of this approach for delaying or mitigating AD progression.

## 1. Introduction

Alzheimer’s disease (AD) is a progressive neurodegenerative disorder that primarily affects older adults and is the leading cause of dementia worldwide (Collaborators, 2022; Western et al., 2024). While the exact causes of AD are still not fully understood, genetic mutations play a significant role, especially in early-onset familial forms of the disease. Three rare single-gene variants, Amyloid Precursor Protein (APP), Presenilin1 (PSEN1), and Presenilin2 (PSEN2), are known to be responsible for these familial cases. APP encodes the amyloid-β precursor protein, which, when cleaved by enzymes such as β-secretase and γ-secretase, generates amyloid-β (Aβ). These amyloid-β peptides aggregate into plaques, a pathological feature found in the brains of AD patients. Despite decades of research into the pathophysiology of AD, the precise molecular mechanisms driving the disease remain unclear, and there is still no effective and safe treatment to slow or halt its progression (Bartoletti-Stella et al., 2018; Uddin et al., 2020). In addition to plaques, neuroinflammation plays a critical role in the development of AD, with elevated pro-inflammatory cytokines, reduced anti-inflammatory responses, and heightened glial cell reactivity. Lipids, particularly fatty acids and their derivatives, have long been recognized for their immunomodulatory effects, capable of triggering either pro-inflammatory or anti-inflammatory responses depending on the lipid class and the target cell (Daynes & Jones, 2002; Leng & Edison, 2021).

As one of the critical aspects of AD pathogenesis, altered lipid profiles and lipid accumulation have been found in both animal models and postmortem human AD brains (Munkacsy et al., 2019; Ralhan, Chang, Lippincott-Schwartz, & Ioannou, 2021). Several lipid-related genes have been identified as genetic risk factors for AD, including Apolipoprotein E (ApoE), ABCA1 (ATP-Binding Cassette Transporter A1), PICALM (Phosphatidylinositol-binding clathrin assembly protein), and TREM2 (Triggering receptor expressed on myeloid cells 2) (Martens et al., 2022; Picard et al., 2018). Notably, the ApoE 4 allele is a well-established genetic risk factor for late-onset AD, with ApoE playing a crucial role in lipid transport and neuronal integrity (Chen et al., 2024). The upregulation of enzymes like fatty acid synthase (FAS) and acetyl-CoA carboxylase (ACC) in AD models suggests disrupted lipid synthesis, with palmitic acid (C16) accumulation near amyloid plaques, highlighting lipid pathways as potential therapeutic targets in AD (Ates, Goldberg, Currais, & Maher, 2020; Daugherty et al., 2017). In addition to neurons, astrocytes are a diverse group of neural cells that primarily degrade fatty acids in the brain. Their metabolic role in AD and neurodegeneration is beginning to be understood, with recent studies suggesting that impaired astrocytic fatty acid degradation may contribute to lipid dysregulation in the brain (Daugherty et al., 2017). This dysfunction in astrocytes could represent a key mechanism driving the pathophysiology of AD. Both microglia and astrocytes express enzymes for β-oxidation, with fatty acid metabolism influencing microglial phenotype and inflammation. Inhibition of β-oxidation in macrophages and microglia exacerbates inflammation, while enhancing fatty acid oxidation reduces lipid-induced inflammation. Activation of mitochondrial function in microglia helps alleviate neuroinflammation and promotes amyloid-β clearance (B. Liu et al., 2019; Namgaladze & Brune, 2016; Song & Suk, 2017). In transgenic mice expressing a mutant APP, there was an increase in *Aβ42* deposits in the brain along with elevated plasma cholesterol levels, further supporting the involvement of disrupted lipid metabolism in AD pathogenesis (Refolo et al., 2000; Shie, Jin, Cook, Leverenz, & LeBoeuf, 2002). Elevated levels of lysophospholipids and ceramides have been detected around Aβ plaques in the human AD brain. Similarly, in *App^NLG-F^* knock-in mice, an age-dependent increase in lysophospholipids and bisphosphates has been observed near Aβ plaques (Huang et al., 2024). Despite these findings, the precise role of lipid metabolism in AD pathology remains underexplored. Understanding how lipid dysregulation contributes to AD could open new avenues for therapeutic intervention.

One of the key enzymes involved in lipid metabolism is diacylglycerol O-acyltransferase 2 (Dgat2), which catalyzes the final step in triglyceride synthesis (Adamovich et al., 2014). Interestingly, Dgat2 has been implicated in both lipid metabolism and the regulation of circadian rhythms, suggesting that its role extends beyond simple lipid storage and may influence the timing of metabolic processes (Livelo et al., 2023; Livelo, Guo, Madhanagopal, Morrow, & Melkani, 2025; Moraes et al., 2024). Dgat2 levels are increased in human AD brains and a mouse AD model (Prakash et al., 2023), with snRNAseq demonstrating its increased expression in excitatory neurons (Brase et al., 2023). Inhibition of its activity in microglia improved microglial uptake of Aβ (Prakash et al., 2025) treating 5xFAD mice for 1 week with a proteasome activator that can degrade Dgat2 in the brain decreased APP in the subiculum (Prakash et al., 2025). Although whether the proteasome activator only degrades microglia Dgat2 is unclear, whether memory is improved by Dgat2 degradation is unknown However, the precise mechanisms by which Dgat2 contributes to AD, particularly in the context of APP mutation and lipid dysmetabolism, remain clear. Circadian rhythms regulate various physiological processes, including sleep, cognition, and metabolism. In AD, circadian rhythms are often disrupted, leading to sleep disturbances and cognitive decline. Interestingly, circadian rhythms influence lipid metabolism, and disruptions in circadian timing can lead to metabolic dysfunction, including altered lipid storage and degradation (Austad et al., 2022; Hardin & Panda, 2013; Panda, Hogenesch, & Kay, 2002). Furthermore, our previous work has shown that modulating lipid metabolism function in *Drosophila melanogaster* (commonly known as the fruit fly), a model organism for studying human aging and pathophysiology (Moraes et al., 2024; Villanueva et al., 2019). We also observed that ApoE induces lipid accumulation in the brain, with ApoE4 uniquely causing this accumulation independently of Dgat2, while ApoE2 and ApoE3 require Dgat2 for lipid buildup (Moraes et al., 2024). These findings highlight the importance of lipid metabolism in AD and suggest that targeting lipid metabolism pathways could mitigate ApoE-related metabolic dysfunction in AD. These findings suggest a strong connection between lipid metabolism and AD pathology (Moraes et al., 2024).

In this study, we use *Drosophila* and *mouse* models of AD to investigate the effect of *APP* and *Aβ42* with and without Dgat2 modulation (*Drosophila*) and with young and old (Mouse) on AD-related phenotypes. Panneuronal, glial, and mushroom body-specific expression of *UAS-App^NLG^, UAS-Aβ42* induced lipid dysmetabolism and neurodegeneration in flies. Our innovative findings in *Drosophila* models of *App^NLG^* mutations and *Aβ42* reveal progressive locomotor impairments, sleep-circadian disturbances, memory deficits, lipid accumulation, and neuroinflammation. Knockdown of Dgat2 improved inflammatory and metabolic gene expression, enhanced locomotor function, and improved sleep activity and circadian rhythms, suggesting the potential benefits of modulating lipid metabolism through Dgat2. For the mouse study, we use *App^NLG-F^* knock-in mice with elevated Aβ pathogenic levels due to the combined effects of three mutations associated with familial AD (Yokoyama, Kobayashi, Tatsumi, & Tomita, 2022). These mice develop amyloid plaques, neuroinflammation, synaptic loss, and cognitive deficits, providing a robust model to study the progression of AD. In this model, we have shown that lipid accumulation and neuroinflammation occur. These findings suggest that targeting Dgat2 may offer a potential therapeutic strategy for AD, emphasizing the conserved impact of lipid metabolism across species.

## 2. Results

A brief outline of our study is shown in **Fig.1A** to reveal the involvement of β-amyloid in AD progression using *Drosophila* and mouse models. As indicated in the method section, using *Drosophila* models, we have tested the impact of *UAS-App^NLG^ and UAS-Aβ42* upon their expression in panneuronal (Elav), glial (GLaz), and mushroom body (Ok-107) drivers. The outline also demonstrated the use of *Dgat2 knockdown (KD)* with these drivers. Geotaxis assays assessed locomotor performance, and sleep-circadian activity was conducted using the DAM system to investigate the impact of APP mutation and Dgat2 modulation. As previously reported, olfactory aversion training was performed to test memory upon expression of APP mutants and Dgat2 modulations. Gene expression analyses of lipid metabolism, neuro-inflammation, and AD risk genes were conducted. All experiments included appropriate controls (W^1118^ and GFP) at two different ages **(Fig. 1A)**. In the *App^NLG-F^*mouse model of AD, along with wild-type control mice, we examined lipid accumulation and glial activation by immunofluorescence. Additionally, qPCR was performed to assess the expression of genes involved in lipid metabolism, neuroinflammation, and AD risks **(Fig. 1A)**.

**Figure 1.**
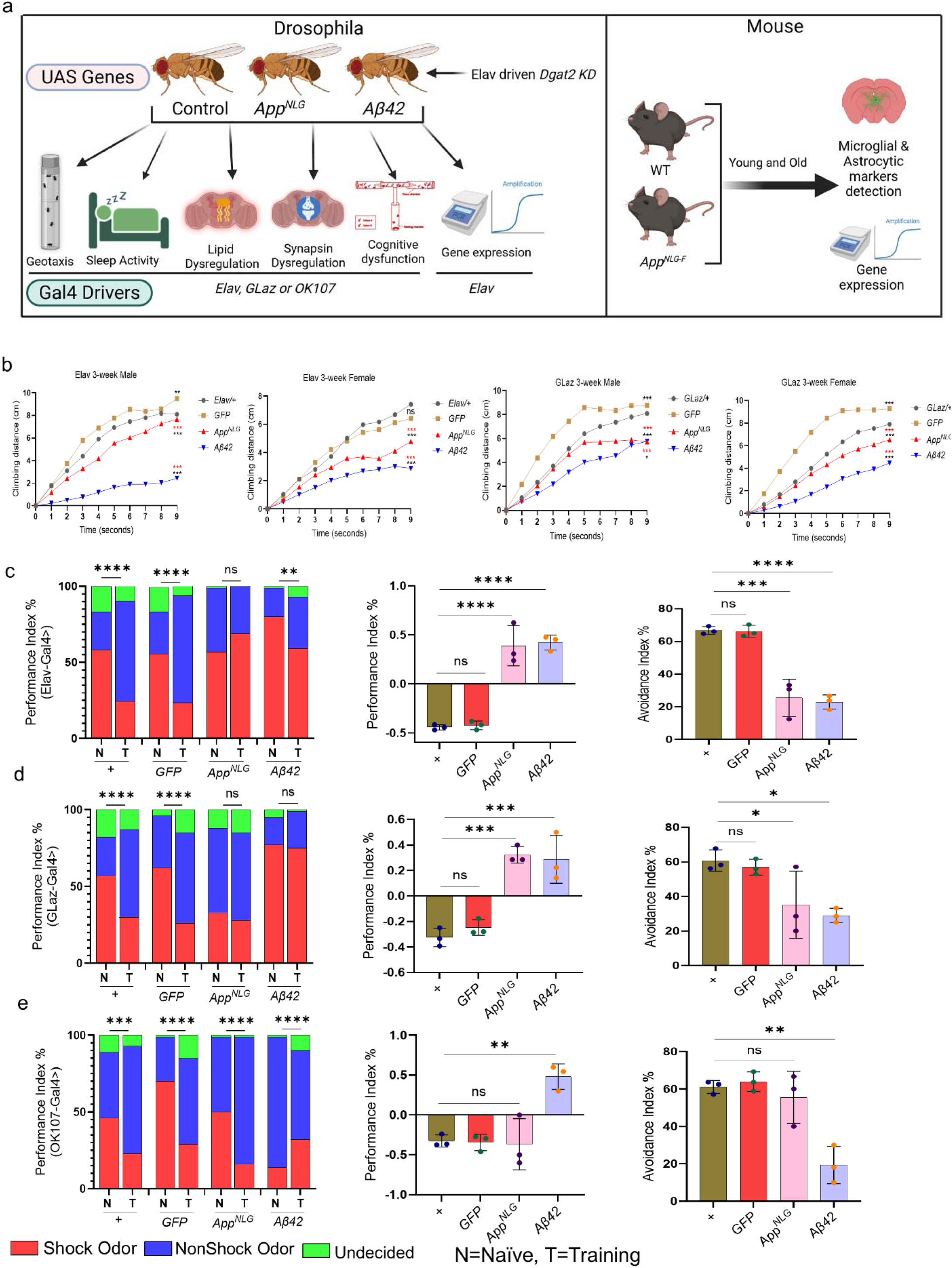
Panneuronal and glial-specific expression of *App^NLG^ and Aβ42* led to compromised locomotor and cognitive performance. (a) Flow chart depicting the experimental setup. Each UAS gene was crossed with the region-specific Gal-4 drivers. To explore the contribution of Dgat2 in *Aβ42 and App* accumulation, we performed Elav-driven *Dgat2 KD* in *Drosophila* and used a young/old *App^NLG-F^*mouse model. (b) Negative geotaxis assay of Elav and GLaz drove A*pp^NLG^ and Aβ42* models in 3-week-old males and females (red asterisk denotes significance with GFP, while black asterisks, ns, denotes significance with Elav/+ and/ or GLaz/+). (c-e, left panels only) *App^NLG^ and Aβ42* models show deficits in olfactory aversion training in 3-week-old male flies in Elav (c) Glaz (d), and OK107 (e), driven models; data combined from triplicate assays. The significance was based on the chi-square test. Values are shown as the percentage of flies in each chamber at the end of the assay. (c-e, middle panels only) The difference between fly decisions, in 3-week-old male flies in Elav (c), Glaz (d), and OK107 (e). (c-e, right panels only), flies avoiding shock odor in Elav (c) Glaz (d), and OK107 (e), driven *App^NLG^ and Aβ42* models. Data = mean ± SD. One-way ANOVA with Tukey’s multiple comparisons test was performed (Left and right panels only). Two-way ANOVA with Tukey’s multiple comparisons test was performed for geotaxis, n=3 replicates per group, and each group had 10 flies. \**P*<0.05, \*\**P*<0.01 and \*\*\**P*<0.001. n.s., not significant (an asterisk denotes significance for the average of all three replicates). Raw data and *P values* are provided in the source data.

### 2.1. Panneuronal, glial, or mushroom-body specific expression of App^NLG^ and Aβ42 led to compromised locomotor and/or cognitive performance

As shown in **Fig.1B**, distinct differences in compromised locomotor performance were observed upon expression of *App^NLG^* and Aβ*42,* with more pronounced impairments in Aβ42-expressing flies and more modest effects in *App^NLG^* females, compared to age-matched controls in 3-week-old flies. Similar trends were seen in male flies with Elav-*Aβ42,* whereas male flies expressing *the App^NLG^* construct showed no significant changes in the climbing ability regardless of the driver used. These results suggest that the geotaxis phenotype is driver-specific and impacts locomotor performance more prominently in females than males, whether through panneuronal or glial expression **(Fig.1B).** To investigate cognitive performance, we subjected wild-type, *App^NLG,^* and *Aβ42* flies to olfactory aversive conditioning, which test their ability to associate odors with reinforcers **(Fig.1C-E)**. 3-week-old female flies expressing *App^NLG^* and *Aβ42* driven by Elav and GLaz resulted in significantly impaired cognitive performance **(Fig. 1C, D left panels).** Expression of these AD-linked genes in the mushroom body using the OK-107 driver led to even greater deficits in olfactory aversion training **(Fig. 1E, left panel).**

Additionally, we performed two olfactory aversion analyses measuring both odors (measured by the odor avoidance index) and shock acuity (measured by the electric shock avoidance index) **(Fig. 1D-E, middle and right panels, respectively).** Both approaches revealed similar conclusions. These findings indicate that the expression of AD-linked genes in mushroom bodies has a more profound impact on memory compared to panneuronal and glial expression, highlighting the critical role of these brain regions in olfactory learning. This suggests that different brain regions contribute to cognitive dysfunction in AD models, offering new insights into the mechanisms underlying AD-related memory impairment.

### 2.2. Compromised sleep-circadian activity in flies with panneuronal and glial-specific expression of Aβ42 and in aged glial-specific expression of App^NLG^

Compromised circadian activity was observed in flies with panneuronal and glial-specific expression of *Aβ42* and in aged glial-specific expression of *App^NLG^*. Various sleep and circadian parameters, including day, night, and total sleep duration, circadian activity, sleep quality, bout length, and bout number, were analyzed in 3 and 7-week-old male flies with panneuronal and glial-specific expression of *App^NLG^* and *Aβ42* **(Figure 2a-f and Figure S1a-d, Figure S2a-f).** These data were compared to controls to evaluate the impact of AD mutations on sleep behavior. As shown in **Figure 2a**, neuronal-specific expression of *Aβ42* led to a significant increase in overall sleep time compared to controls in 3-week-old flies, particularly during the night **(Figure 2a).** The increased sleep was accompanied by lower sleep activity in *Aβ42* **(Figure 2c).** Although a similar trend was observed with *App^NLG,^* changes were not statistically significant. These findings suggest that while the panneuronal-specific expression of *Aβ42* led to more sleep, the quality of sleep was compromised due to lower sleep bout number and higher bout length **(Figure S1a, b).**

**Figure 2.**
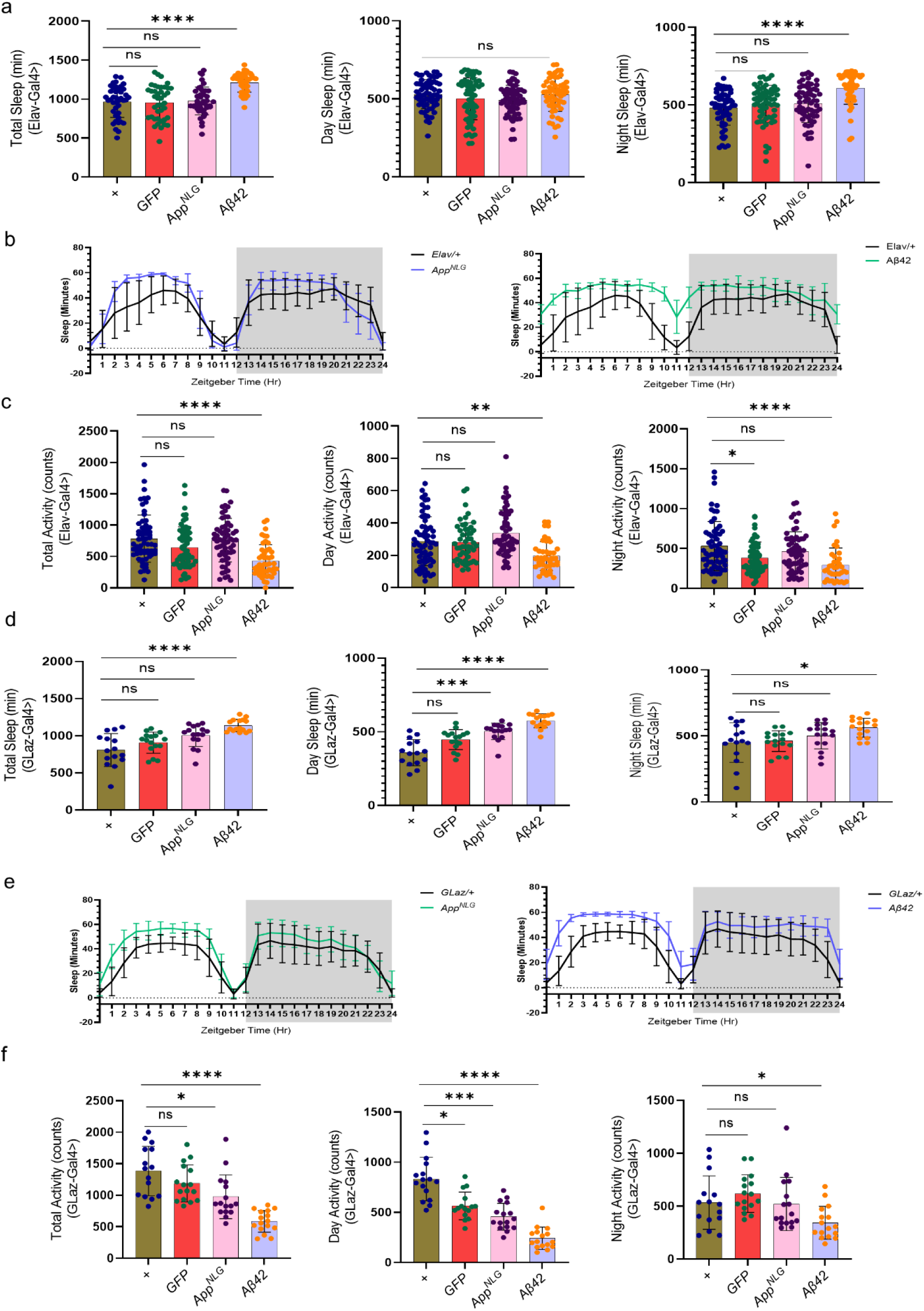
Panneuronal and glial-specific expression of Aβ*42* has compromised circadian activity and sleep quality compared to *App^NLG,^* which worsens with age. (a) Total, day and night sleep (in minutes) in Elav-driven *App^NLG^* and Aβ*42 m*odels. (b) Sleep profiles at different zeitgeber time (hr.) in Elav-driven *App^NLG^* and Aβ*42* models only compared to their respective controls. (c) Total, day and night sleep activity (counts) in Elav-driven *App^NLG^*and Aβ*42* models. (d) Total, day and night sleep in (in minutes) in GLaz-driven *App^NLG^* and Aβ*42* models. (e) Sleep profiles at different zeitgeber time (hr.) in GLaz-driven *App^NLG^* and Aβ*42* models only compared to their respective controls. (f) Total, day and night sleep activity in GLaz-driven *App^NLG^*and Aβ*42* models. All experiments were performed in 3-week-old males only. Data = mean ± SD. Non-parametric One-way ANOVA with multiple comparisons, done with the Kruskal-Walli’s test, was performed for sleep activity and sleep fragmentation. Each dot represents the number of flies. \**P*<0.05, \*\**P*<0.01 and \*\*\**P*<0.001. n.s., not significant (an asterisk denotes significance for the average of all three replicates). Raw data and *P values* are provided in the source data.

As shown in **Figure 2d-f and Figure S1d, e,** flies with glial-specific expression *App^NLG^* and *Aβ42* genes exhibited significantly increased sleep, especially in those expressing *Aβ42.* This increase was mainly due to longer sleep during the night, as illustrated in the 24-hour sleep profile, which shows sleep minutes/ hours **(Figure 2e).**

This increase in sleep was associated with lower sleep activity during the day and nighttime in both *App^NLG^*and *Aβ42* **(Figure 2f)** and increased bout length in both *App^NLG^* and *Aβ42* **(Figure S1c, d),** suggesting that while the total sleep duration increased, the quality of sleep was negatively impacted. The increased sleep duration was thus accompanied by poor sleep quality, as indicated by reduced activity.

Additionally, panneuronal expression of *Aβ42* led to a decrease in the total number of sleep bouts, especially at night, suggesting fewer, longer sleep periods **(Figure S1a, b).** In contrast, glial-specific expression of *App^NLG^* and *Aβ42* caused an increase in the number of sleep bouts during the day, with longer day bout lengths, indicating a distinct pattern of sleep fragmentation **(Figure S1c, d).** In 7-week-old flies, sleep disturbances persisted and worsened, indicating that these disruptions are not transient but progressively worsen with age (**Figure S2a-f).** These findings suggest that sleep disturbances are a lasting feature of AD models.

### 2.3. Panneuronal and glial-specific expression of App^NLG^ and Aβ42 led to increased lipid accumulation and reduced synapsin levels

Our study revealed a significant increase in lipid accumulation, as indicated by green positive cells (Lipid spot) in both the *App^NLG^* a*nd Aβ42* flies, compared to their respective control groups at 3 weeks **(Figure 3a-f).** Synapsin staining intensity was significantly lower in these flies compared to controls, indicating impaired synaptic function **(Figure 3d, f).** This reduction was observed in both panneuronal and glial-driven *App^NLG^ and Aβ42* flies. At 7 weeks, lipid accumulation remained elevated in both the *App^NLG^*a*nd Aβ42* flies in both panneuronal and glial drivers. While synapsin level remains unchanged in panneuronal-driven flies, but was reduced in glial-driven *App^NLG^*model, suggesting a stabilization or compensatory adaptation in synaptic function (**Figure S3-f**). These findings indicate an early phase of synaptic damage followed by a potential plateau in synaptic decline, with persistent lipid and amyloid changes pointing to ongoing pathophysiological processes. Overall, the results suggest that lipid dysregulation and synapsin impairment are strongly linked in AD progression, with the *App^NLG^* model potentially reflecting more closely the synaptic dysfunction observed in AD, which may contribute to cognitive and sleep disturbances.

**Figure 3.**
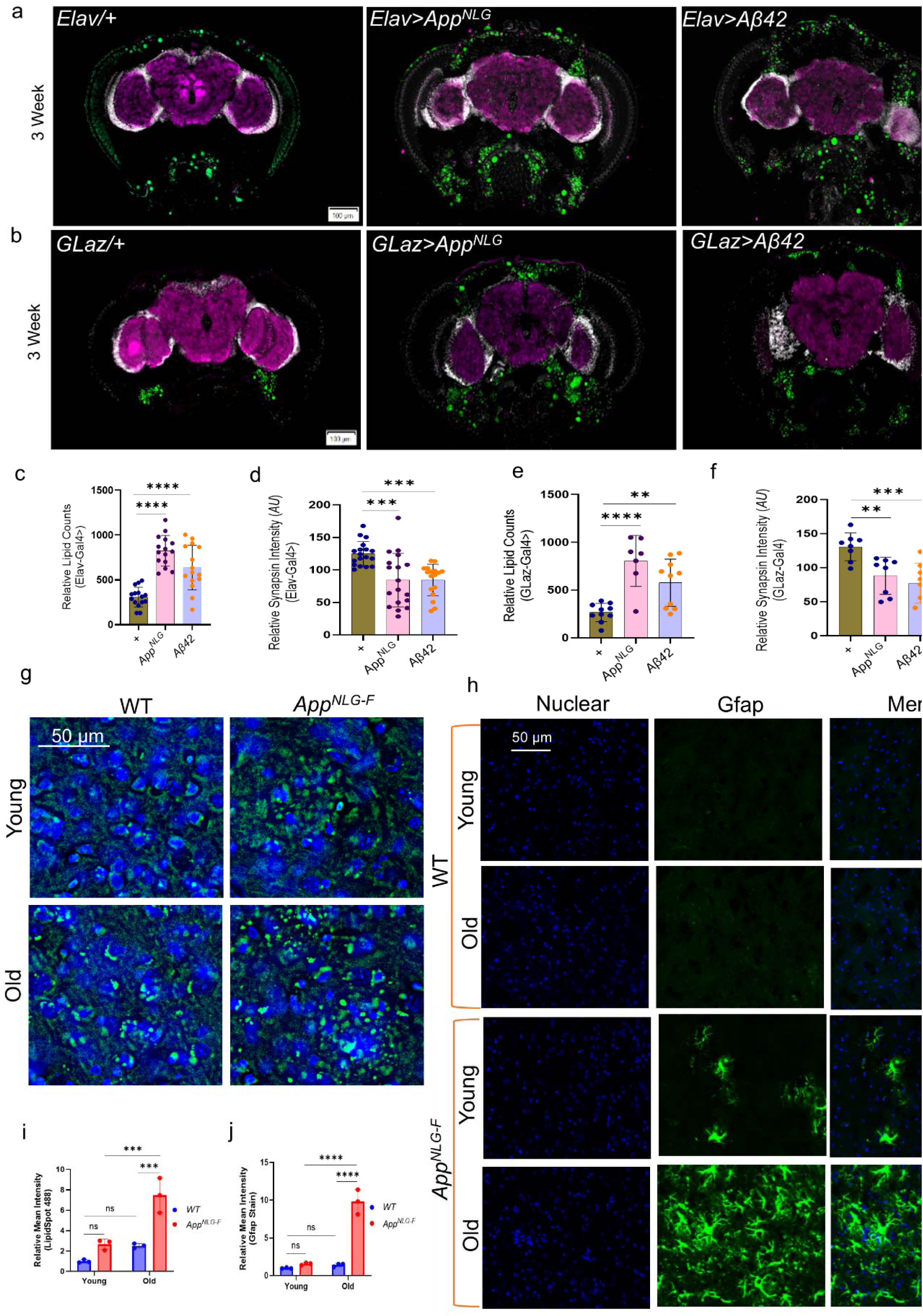
*App^NLG^ and Aβ42* induce lipid accumulation and synapsin reduction in Drosophila, while *App^NLG-F^*mice show cortical lipid build-up and astrocyte activation. (a, b) Representative image showing the expression of lipid accumulation (green, Lipid spot); synaptic loss (purple, anti SYNORF1), a marker of neurodegeneration; and DAPI (white), in the brains of Elav and GLaz**-**driven *App^NLG^ and Aβ42* flies. (c-f) Quantification of the expression level of relative lipid counts (c, e) and synapsin intensity (d, f) in Elav and GLaz-driven *App^NLG^ and Aβ42* flies. All experiments were performed in 3-week-old flies. (g) Representative images showing lipid spot intensity (green), indicating lipid accumulation; and DAPI (blue), in the mouse brains of young and old wild type (WT) and *App^NLG-F^*models. (h) Representative images showing the immunostaining of Gfap, a marker of astrocyte activation; and DAPI (blue), in the mouse brains of young and old wild type (WT) and *App^NLG-F^* models. (i, j) Quantification of the intensity of lipid spots and relative mean intensity of Gfap immunostaining. Fold changes of fluorescence intensity were calculated relative to controls. Data = mean ± SD. One-way ANOVA with Tukey’s multiple comparisons test was performed for flies’ data, with 5-6 flies per group. Two-way ANOVA with multiple comparisons done with Uncorrected Fisher’s LSD test, was performed for mouse data, with n=3 mice per group. \**P*<0.05, \*\**P*<0.01 and \*\*\**P*<0.001. n.s., not significant (an asterisk denotes significance for the average of all three replicates). Raw data and *P values* are provided in the source data.

### 2.4. App^NLG-F^ mice exhibit an age-dependent increase in lipid accumulation and glial activation

We performed immunostaining in the cortex (Ctx) region of the *App^NLG-F^* mice to assess the level of lipid spot staining and Gfap immunoreactivity, respectively, at different ages (young and old mice). In 15-month-old *App^NLG-F^* mice, we observed a notable increase in lipid accumulation, as indicated by enhanced lipid spot intensity **(Figure 3g, i)**. Additionally, Gfap level, which marks astrocytic activation, was significantly higher in 15-month-old *App^NLG-F^* compared to age-matched wild-type (WT) controls **(Figure 3h, j).**

We also examined Nile red and Dgat2 staining in the Ctx **(Figure S3g-j).** Nile red staining significantly increased in old *App^NLG-F^*mice, while Dgat2 staining showed no change in intensity, although Dgat2 aggregates were observed in the hippocampal CA1 region and cerebellum **(Figure S4a-d).** Furthermore, we evaluated Adrp, a protein that binds lipid droplets, and found no significant changes in its levels in the cortex or hippocampal (CA1) regions **(Figure S5a-d).** Immunostaining for Gfap and Iba1 in the CA1 region **(Figure S7a-d)** revealed a significant increase in Gfap-positive cells in 15-month-old *App^NLG-F^* mice. Iba1-positive cells, marking microglial activation, were also significantly higher in both young and old *App^NLG-F^* mice compared to WT controls. Notably, aging did not affect Iba1 expression in WT mice, but a dramatic increase in Iba1 was observed in 15-month-old *App^NLG-F^* mice, both in the CA1 region **(Figure S6a, b)** and in the cortex (**Figure 4a, b)**. These findings suggest that aging in *App^NLG-F^* mice is associated with increased lipid accumulation, astrocytic activation, and microglial changes, consistent with age-related neuroinflammation in AD models.

**Figure 4.**
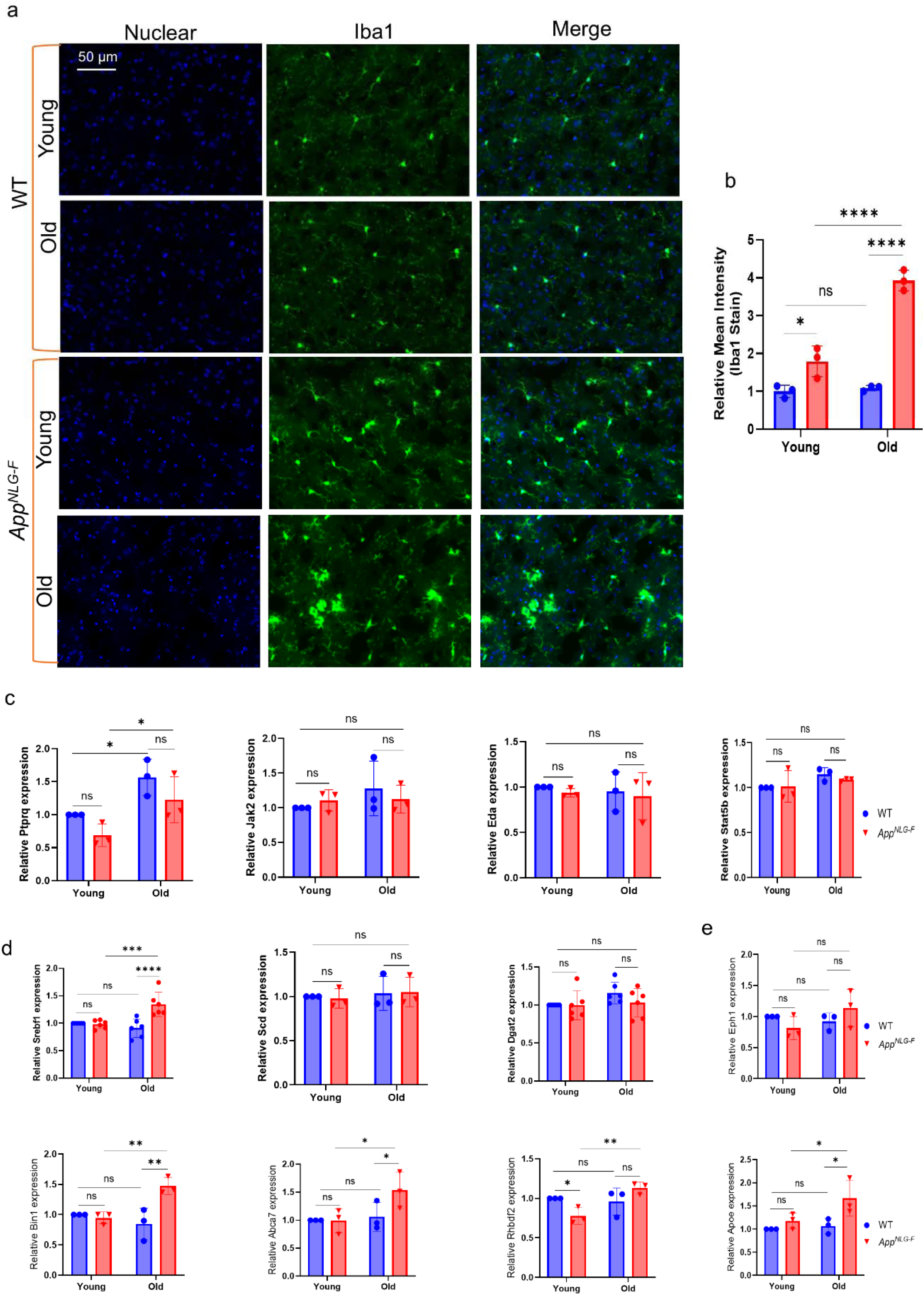
Age-dependent increase in neuroinflammation and AD risk gene expression in a mouse model of *App^NLG-F^*. (a) Immunofluorescence staining of Iba1 protein in microglia within the cortex region of the brain of WT and *App^NLG-F^* mice at young and old age. (b) Quantification of the Iba1 immunostaining intensity in (a); n=3 mice per group. (c-e) Quantitative PCR quantification of the relative expression level of inflammatory genes, Ptprq, Jak2, Eda, and Stat5b (c), metabolic genes, srebf1, scd, and Dgat2 (d), and AD risk genes, Bin1, Abca7, Rhbdf2, and Apoe (e), in young and old WT and *App^NLG-F^* mice. Data = mean ± SD. Two-way ANOVA with multiple comparisons, done with Uncorrected Fisher’s LSD test, was performed. \**P*<0.05, \*\**P*<0.01 and \*\*\**P*<0.001. n.s., not significant (an asterisk denotes significance for the average of all three replicates). Raw data and *P values* are provided in the source data.

### 2.5. Age-dependent increase in AD risk gene expression in the App^NLG-F^ mouse model

In the *App^NLG-**F**^* mice, we observed a significant age-dependent increase in Iba1, which is a marker of microglia activation **(Figure 4a, b and Figure S6b, d).** This rise in Iba1 suggests that microglia become more activated with age, possibly due to the amyloid plaques in the brain. Microglia initially respond to plaques and other damage by becoming activated to clear harmful debris, but chronic activation over time can lead to neuroinflammation. This persistent activation can reduce the efficiency of microglial function, contributing to neuronal damage and the progression of neurodegenerative diseases like Alzheimer’s. To explore this further, we examined the expression of key inflammatory genes. Notably, protein tyrosine phosphatase receptor type Q (Ptprq), showed a mark upregulation with age **(Figure 4c)**, supporting the idea of chronic, age-related inflammation. However, other inflammatory markers, such as Janus kinase2 (Jak2), ectodysplasin (Eda), and signal transducer and activator of transcription 5B (Stat5b) did not show significant change, suggesting a more selective inflammatory response. In addition to neuroinflammation, we assessed the expression of metabolic genes. Notably, Srebf1 expression was significantly increased with age, while no significant changes were observed in other metabolic genes, such as stearoyl-CoA desaturase (Scd) and Dgat2, indicating a possible alteration in lipid metabolism (**Figure 4d)**.

Further investigation of AD risk genes revealed an age-dependent increase in the expression of Bridging integrator 1 (Bin1) and ATP binding cassette subfamily A member 7 (Abca7), both of which are involved in amyloid processing and lipid transport **(Figure 4e).** This upregulation suggests a potential link to increased amyloid burden or alterations in lipid metabolism, both of which are key features of Alzheimer’s pathology. Additionally, we observed elevated expression of rhomboid family-like 2 (Rhbdf2) and Apoe genes that are also associated with neuroinflammation and lipid homeostasis **(Figure. 4e),** and the age-dependent increases of Bin1 and Abca7 are only shown in AD models but not in WT mice.

Taken together, the increased expression of Iba1, Ptprq, and several AD risk genes with age highlights the progressive activation of neuroinflammatory pathways and suggests that lipid metabolism and inflammatory responses are central contributors to the neurodegenerative processes in the *App^NLG-F^*mouse model. This growing evidence reinforces the idea that neuroinflammation and the dysregulation of AD-related pathways progressively exacerbate pathology as the mice age.

### 2.6. App^NLG^ and Aβ42-linked locomotor and cognitive dysfunction was improved with Dgat2 KD

We assessed geotaxis, and cognitive function, in 3-week-old flies to investigate the effects of *App^NLG^* and *Aβ42* on the panneuronal expression of *Dgat2KD*. First, a geotaxis assay was used to assess locomotor performance. At 3 weeks of age, significant differences in locomotor performance were observed between flies expressing the *App^NLG^* gene and their respective controls. Male flies with panneuronal *Dgat2 KD* displayed improved climbing ability when expressing *App^NLG^ or Aβ42, w*hile female flies show similar trends, though less significantly **(Figure 5a).** These results suggest that Dgat2 KD may protect locomotor performance.

**Figure 5.**
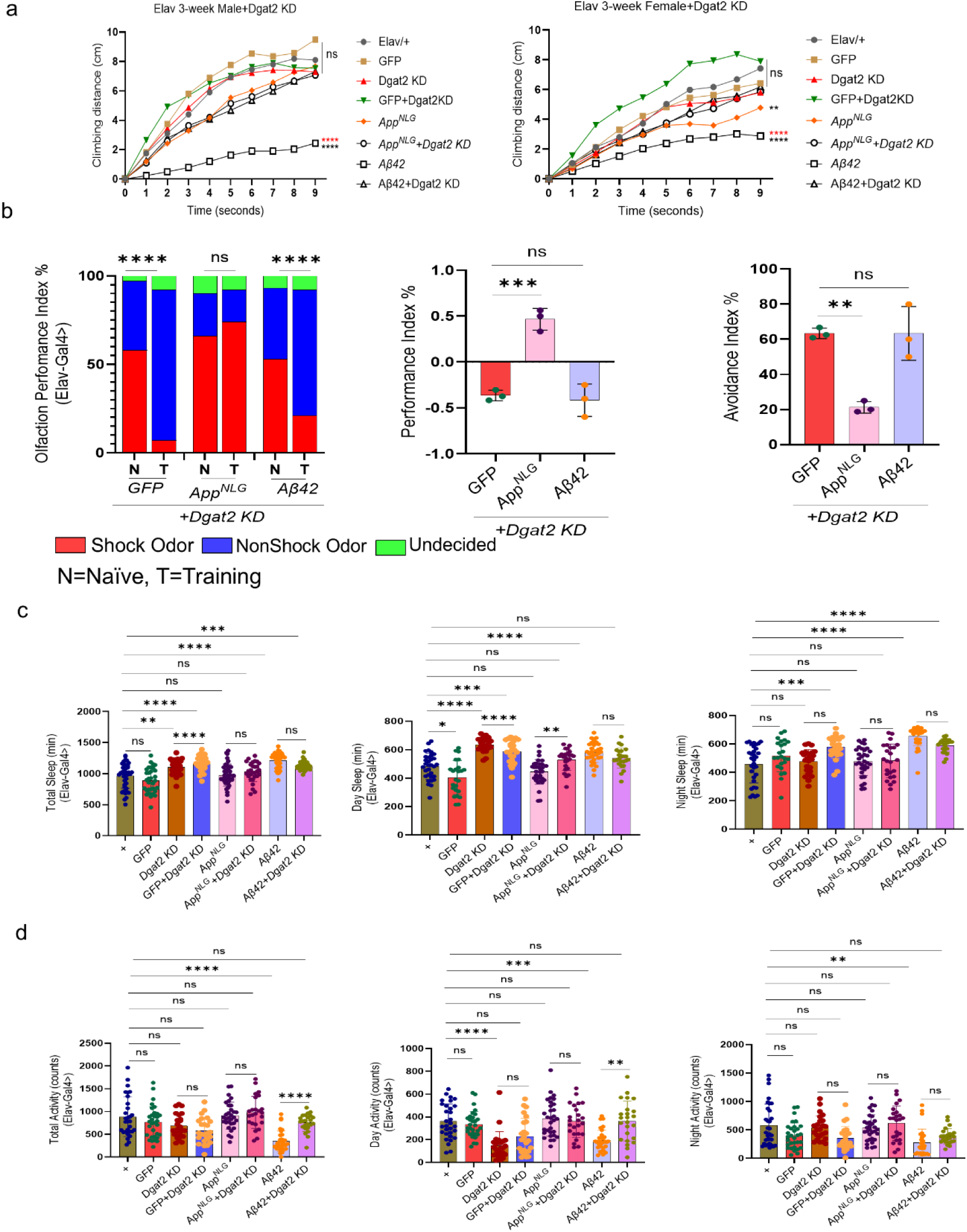
*Dgat2 KD* enhances locomotor and cognitive performance, restores sleep activity, and reduces fragmentation linked to *App^NLG^ and Aβ42* expression. (a) Negative geotaxis assay of Elav-driven *Dgat2 KD* in A*pp^NLG^ and Aβ42* models in 3-week male and female (red asterisk denotes significance with *Aβ42+Dgat2 KD* and/or *App^NLG^ +Dgat2 KD while* black asterisks, ns, denotes significance with *Elav/+*). (b, left panels only) *Dgat2 KD* overcame deficits in olfactory aversion training in Elav-driven 3-week-old male *App^NLG^ and Aβ42* models; data combined from triplicate assays. The significance was based on the chi-square test. Values are shown as the percentage of flies in each chamber at the end of the assay. (b, middle panels) the difference between fly decisions in Elav-driven *Dgat2 KD* in 3-week-old male *App^NLG^ and Aβ42* models. (b, right panels), flies avoiding shock odor in Elav-driven *Dgat2 KD* in 3-week-old male *App^NLG^ and Aβ42* models. (c) Total, day and night sleep (in minutes) in Elav-driven *Dgat KD* in *App^NLG^ and Aβ42* models. (d) Total, day and night sleep activity (in counts) in Elav-driven Dgat KD in *App^NLG^ and Aβ42* models. Data=mean ± SD. One-way ANOVA with Tukey’s multiple comparisons test was performed (Left and right panels only). Two-way ANOVA with Tukey’s multiple comparisons test was performed for geotaxis, n=3 replicates per group, and each group had 10 flies. Non-parametric One-way ANOVA with multiple comparisons, done with the Kruskal-Wallis test, was performed for sleep activity and sleep fragmentation. \**P*<0.05, \*\**P*<0.01 and \*\*\**P*<0.001. n.s., not significant (an asterisk denotes significance for the average of all three replicates). Raw data and *P values* are provided in the source data.

We next evaluated if cognitive performance was altered in *Dgat KD*. Male flies expressing *GFP, Aβ42,* and *App^NLG^* exhibited memory impairment, which was rescued when Dgat2 was KD **(Figure 5B, left panel)**. Further analysis of odor avoidance using the performance index **(Fig. 1c-e)** supported these findings, reinforcing the idea that Dgat2 manipulation affects both locomotion and cognition **(Figure 5b, c, middle and right panel).** In conclusion, our results suggest that Dgat2 KD has a protective effect, improving both locomotor and cognitive function in these areas.

### 2.7. Dgat2 KD leads to the restoration of sleep activity linked with the expression of App^NLG^ and Aβ42

We assessed sleep activity in Elav-driven Dgat2 KD and at both 3 and 7 weeks of age to explore the potential links between sleep, cognitive function, and memory performance, which were observed in earlier tests in *App^NLG^* and *Aβ42*. At 3 weeks of age, Dgat2 KD flies expressing *Aβ42* exhibited a more prominent increase in total sleep, including daytime and nighttime sleep compared to *App^NLG^*. Dgat2 KD showed increased sleep timing, indicating a more continuous sleep pattern **(Figure 5c, d).** These suggest that Dgat2 manipulation at 3 weeks drastically restores the sleep pattern of *App^NLG^* and *Aβ42*. Further investigation revealed changes in sleep activity in the *App^NLG^* and *Aβ42* models, such as fewer sleep bouts and increased bout length, particularly in flies with Elav-driven *Dgat2 KD* **(Figure S9).** However, at 7 weeks, sleep behavior in *App^NLG^* and *Aβ42* models with *Dgat2 KD* was largely unchanged compared to the 3-week-old flies **(Figure S10a-d).** This suggests that the impact of Dgat2 manipulation on sleep may be age-dependent, with no significant change in older flies.

### 2.8. Dgat2 knockdown suppressed lipid metabolism, restored synaptic loss, and reduced inflammation associated with App^NLG^ and Aβ42

We assessed immunostaining in Elav-driven *Dgat2 KD* flies expressing *App^NLG^ and Aβ42* at 3 weeks to examine potential changes in lipid metabolism and synaptic function. At this early age, we observed that lipid accumulation, typically elevated in *App^NLG^ and Aβ42* models, was significantly reduced in the presence of *Dgat2 KD.* This suggests that *Dgat2 KD* mitigates abnormal lipid buildup associated with AD models. Interestingly, synapsin levels were notably elevated in these flies **(Figure 6a-c),** indicating improved synaptic function in response to decreased lipid accumulation. At 7 weeks, lipid levels remained reduced in the presence of *Dgat2 KD,* suggesting sustained mitigation of lipid dysregulation. However, synapsin levels were unaffected at 7 weeks **(Figure S10a-c),** indicating that neither the expression of the *App* mutant or β-amyloid nor the continued reduction in lipid accumulation due to Dgat2 knockdown, synapsin levels remain unchanged at this age. This suggests that *Dgat2 KD* has a persistent effect on lipid metabolism, but its impact on synaptic function may be age dependent.

**Figure 6.**
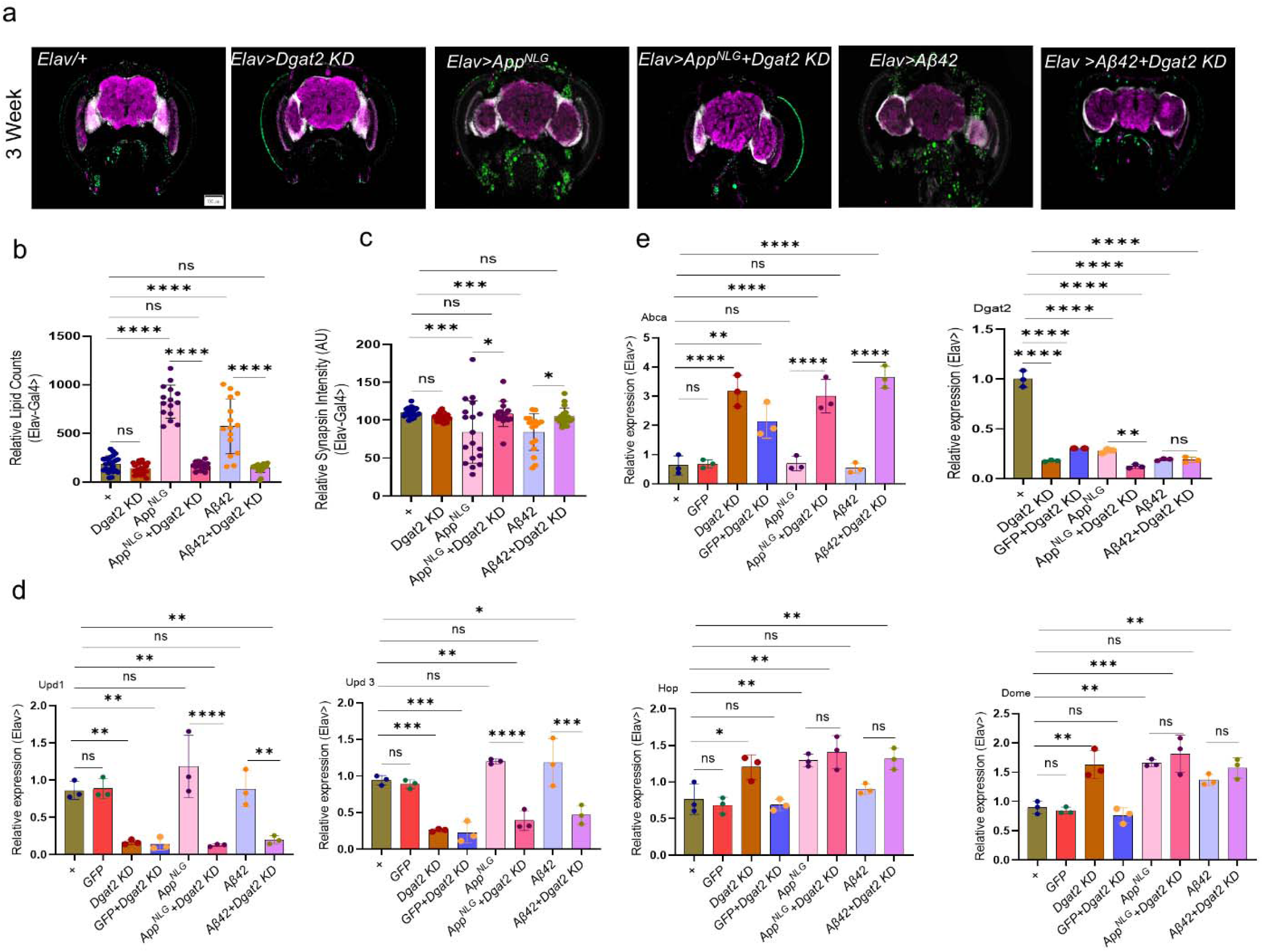
*Dgat2 KD* suppresses lipid metabolism, restores synaptic loss, and reduces inflammation in *App^NLG^ and Aβ42* models of *Drosophila*, while Dgat2 inhibition modulates AD risk genes in *App^NLG-F^* mice. (a) Representative image showing the expression of lipid accumulation (green, Lipid spot 488); synaptic loss (purple, anti SYNORF1); a marker of neurodegeneration, and DAPI (white), in the brains of Elav-driven KD in *App^NLG^ and Aβ42* models. (b, c) Quantification of the expression level of lipid counts and synapsin intensity in Elav-driven *App^NLG^ and Aβ42* models. (d) Relative expression of inflammatory genes, Upd1, Upd3, Hop and Dome in flies. (e) Relative expression of metabolic genes, Abca and Dgat2 in flies. Data=mean ± SD. One-way ANOVA with Tukey’s multiple comparisons test was performed on flies and mouse data. n=3 replicates per group and each group has 8-10 flies. \**P*<0.05, \*\**P*<0.01 and \*\*\**P*<0.001. n.s., not significant (an asterisk denotes significance for the average of all three replicates). Raw data and *P values* are provided in the source data

We further examined neuroinflammatory markers. The levels of primary inflammatory cytokines, unpaired1 (Upd1) and unpaired3 (Upd3), were significantly reduced in *Dgat2 KD* flies, indicating decreased neuroinflammation. However, expression of the JAK/STAT pathway receptors, domeless (Dome) and Hopscotch (Hop), was increased, suggesting that the reduction in inflammation may activate compensatory signaling through this pathway **(Figure 6d).** We also analyzed the expression of metabolic markers *Abca* and *Dgat2*. In both *App^NLG^* and *Aβ42* models, Dgat2 knockdown led to a marked reduction in Dgat2 expression, confirming the efficiency of the knockdown and validating its use in these neurodegenerative backgrounds. At the same time, we observed a consistent increase in *Abca* expression under these conditions. This reciprocal pattern suggests that loss of Dgat2, an enzyme central to triglyceride synthesis and lipid droplet formation, perturbs lipid metabolism and elicits a compensatory upregulation of *Abca*. Together, these findings indicate that Dgat2 downregulation disrupts lipid storage pathways, while Abca upregulation may act to maintain lipid homeostasis, underscoring a critical interplay between lipid synthesis and transport in neurodegeneration (**Figure 6e**)

Overall, we have shown a summary of the behavior, physiological and cytological alterations with the expression of AD-linked genes in *Drosophila* and mouse models. Moreover, the impact of Dgat2 on modulating AD phenotype has also been summarized in **Figure 7**. These findings underscore the intricate connection between amyloid pathology, lipid dysregulation, and neuroinflammation, suggesting that targeting Dgat2, through conserved lipid homeostasis mechanisms across species, may offer a novel and translationally relevant therapeutic approach for Alzheimer’s disease.

**Figure 7.**
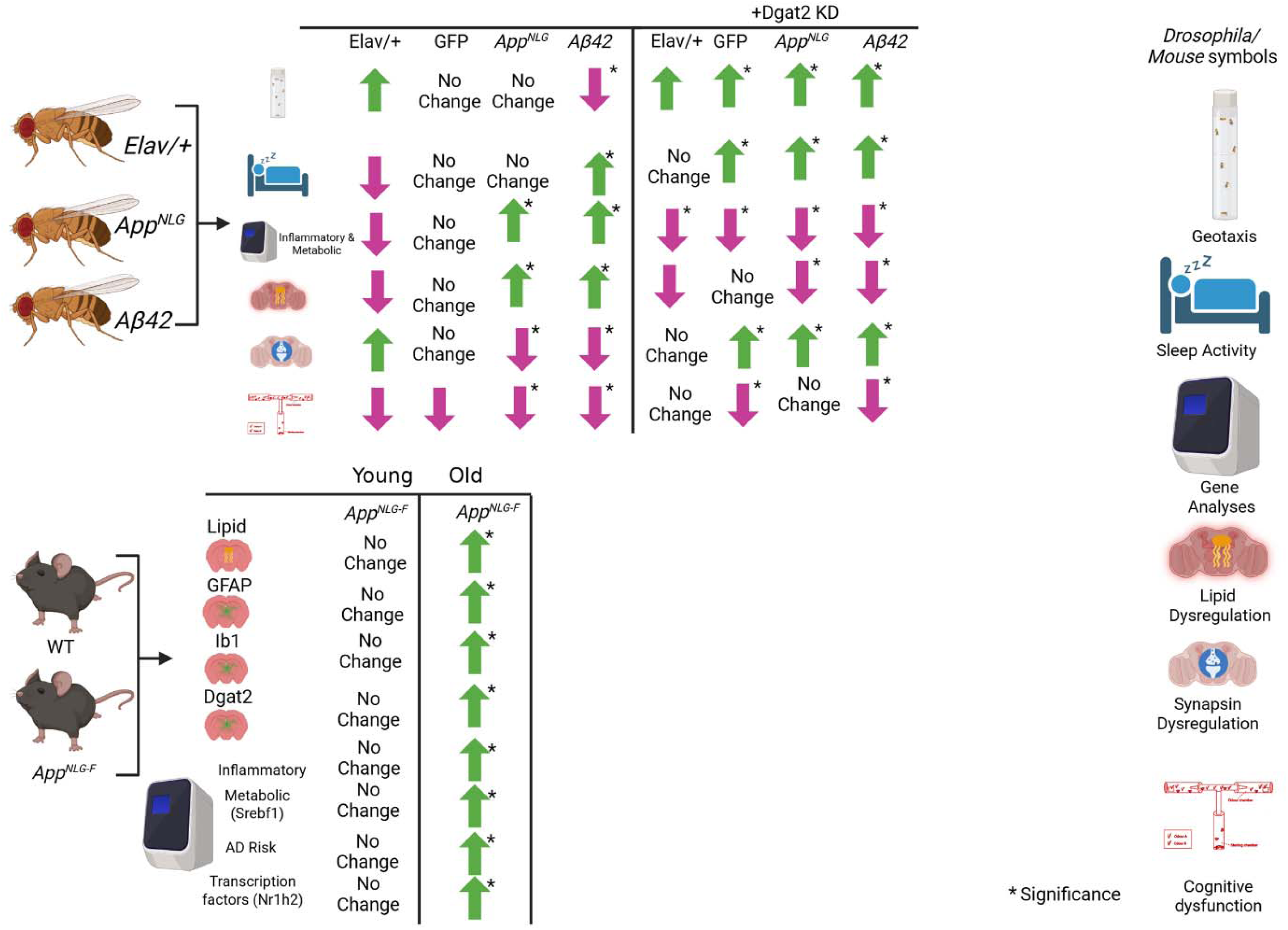
Graphical illustration showed the impacts of panneuronal and glial-specific expression of *App^NLG^ and Aβ42* without and with *Dgat2 KD* in Drosophila and microglial/ astrocytic activation and lipid accumulation in *App^NLG-F^*mice. The top panel result shows that panneuronal and glial-specific expression of *App^NLG^ and Aβ42* led to compromised locomotor and cognitive performance. It also led to enhanced lipid accumulation and reduced synapsin levels, which were further rescued with *Dgat2 KD* (up and down arrows). The bottom panel shows the changes in Gfap, Iba1, Dgat2 stain, inflammatory, and AD risk gene expression in young/old *App^NLG-F^* mice.

## 3. Discussion

AD is characterized by the accumulation of amyloid plaques and tau tangles, leading to cognitive decline and neurodegeneration (Zhang et al., 2021). Genetic models, such as *Drosophila* expressing APP mutations, offer valuable insights into the mechanisms underlying AD (Padmanabhan & Gotz, 2023). This study explored the effects of *App^NLG^*, as well as the aggregation-prone *Aβ42*) on locomotor function, memory, and lipid metabolism in *Drosophila.* Additionally, we examined the role of Dgat2, a key enzyme in lipid metabolism, through its knockdown (Rong et al., 2024). To further establish our findings, we used the *App^NLG-F^* mouse model of AD, allowing for cross-species comparison of lipid profiles, neuroinflammation, and gene expression. Through locomotor assays, cognitive tests, lipid analysis, and gene expression studies, we aimed to uncover how these genetic modifications influence AD pathogenesis and identify potential therapeutic strategies for mitigating AD-associated dysfunction.

Our findings highlight the complex interplay between amyloid pathology, lipid metabolism, and neuroinflammation across the species using the *Drosophila* and mouse AD models. We observed that male *Drosophila* expressing the *Aβ42* mutation exhibited significant locomotor impairments, particularly reduced climbing ability in the geotaxis assay. Interestingly, the decline was more pronounced in the flies with panneuronal or glial-specific expression of *Aβ42* compared to those expressing the *App^NLG^* construct, suggesting that the geotaxis phenotype is mutation-specific **(Figure 1b).** This aligns with previous studies linking amyloid-related motor deficits to the expression of toxic *Aβ* species (Sun, Hu, Wu, Zhou, & Huang, 2022). Cognitive assays also revealed a significant memory impairment, particularly in flies expressing *Aβ42* in the mushroom bodies, a key brain region critical for olfactory learning **(Figure 1c-e)**. This observation underscores the critical role of specific brain regions in cognitive dysfunction in AD, aligning with findings in mammalian models where amyloid pathology in the hippocampus and cortex impairs memory (Polis & Samson, 2024).

Our findings demonstrate significant sleep disruptions in *Drosophila* models expressing the *Aβ42* mutation, particularly in terms of sleep duration and altered sleep patterns. These disruptions were more pronounced compared to the APP mutation model, indicating that the *Aβ42* mutation has a stronger impact on sleep quality. The increase in total sleep time, predominantly during the nighttime **(Figure 2a, b),** aligns with previous research showing that AD models often exhibit prolonged sleep durations, especially at night (Insel, Mohlenhoff, Neylan, Krystal, & Mackin, 2021; Y. Liu et al., 2021). However, our results also revealed a notable decrease in sleep activity (counts), particularly in the Elav- and GLaz-driven models, suggesting that while these flies sleep more, the quality of their sleep is impaired **(Figure 2a-f).** Furthermore, decreased activity levels, particularly in the Elav-driven model, suggested that the sleep changes are closely tied to disruptions in overall activity and circadian rhythms, as seen in both *Drosophila* and mammalian AD models (Duncan et al., 2022; Majcin Dorcikova, Duret, Pottie, & Nagoshi, 2023; Winer et al., 2021). The persistence of sleep disturbances in older flies suggests that sleep dysregulation is a chronic feature of AD pathology, much like the long-term sleep issues observed in AD patients (Bellesi et al., 2017; Minakawa et al., 2017). The differences in sleep fragmentation patterns between the neuronal and glial-specific models highlight the potential role of glial cells in modulating sleep, with glial-specific expression leading to a distinct pattern of increased daytime sleep bouts and longer bout lengths. This is consistent with recent studies implicating glial activation in the disruption of sleep architecture in AD models (Bellesi et al., 2017; Garofalo et al., 2020; Gentry et al., 2022). Collectively, these results reinforce the idea that sleep disturbances, particularly fragmented sleep, are a central feature of AD pathology and may serve as an early indicator of disease progression.

Evidence from our study provides that lipid dysregulation and synaptic impairment play crucial roles in the progression of AD, with distinct differences in the severity of these alterations between the *Aβ42 and App^NLG^* models. Lipid accumulation was significantly elevated in both Drosophila and mouse models of AD, particularly in the *Aβ42* model, which exhibited more pronounced lipid deposition compared to the *App^NLG^* model (**Figure 3a-f**). This aligns with previous research indicating that dysregulated lipid metabolism, particularly the accumulation of lipid droplets, is a key feature of AD pathogenesis (Y. Liu et al., 2021). These lipid imbalances are thought to contribute to cellular stress and dysfunction, particularly in neurons and glial cells, which may exacerbate the cognitive and behavioral deficits seen in AD (Fan et al., 2020; Keeney et al., 2013).

In addition to lipid accumulation, we observed a marked reduction in synapsin expression, a protein crucial for synaptic function **(Figure 3d, f).** This finding is consistent with studies showing that synaptic loss and dysfunction are hallmarks of AD and are often correlated with cognitive decline (Lan et al., 2022; Saroja, Sharma, Hof, & Pereira, 2022). The more pronounced reductions in synapsin in the *Aβ42* model suggest that amyloid-β may have a more severe impact on synaptic integrity than other forms of APP mutations (Piccoli et al., 2022). Transitioning to the *App^NLG-F^*mouse model further corroborates these findings, showing a significant increase in lipid accumulation and glial activation, particularly with aging, suggesting that lipid dysregulation and neuroinflammation are progressive features of AD pathology **(Figure 3g-j, Figure S3g-j, S4, S5).** Aging exacerbates lipid accumulation and astrocyte activation, a key aspect of AD progression (Emre et al., 2021). Elevated GFAP level, indicative of glial activation, further supports the idea that lipid imbalances are coupled with neuroinflammation in the aging brain (Schuitemaker et al., 2012).

Microglial activation and neuroinflammation play pivotal roles in AD progression, as evidenced by the increased Iba1, a marker for microglial activation, in the *App^NLG-F^* mouse model **(Figure 4a, b).** This age-dependent increase of Iba1 suggests that chronic neuroinflammation becomes more pronounced with advancing age, consistent with studies linking microglial activation to neurodegenerative diseases, including AD (Keeney et al., 2013). The increase in Iba1 is consistent with reports indicating that sustained microglial activation is a driver of neurodegeneration (Agrawal & Jha, 2020; Sakuraba, Shintani, Tani, & Noda, 2013). Furthermore, the observed upregulation of Ptprq **(Figure 4c)** corroborates the presence of chronic, age-related inflammation in these mice, consistent with other reports on the role of neuroinflammation in AD pathology (Prendecki et al., 2020). Interestingly, the increased expression of key AD risk genes, including Bin1, Rhbdf2, Apoe and Abca7, which are involved in amyloid processing and lipid metabolism **(Figure 4e),** suggests that amyloid burden or lipid dysmetabolism worsen with age, further implicating lipid pathways in AD progression (De Jager et al., 2014). Furthermore, the elevation of Rhbdf2 and ApoE, both of which are linked to neuroinflammation and lipid homeostasis, underscores the role of disrupted lipid pathways in AD risk (Stone et al., 2009). Collectively, these findings point to a progressive activation of inflammatory and amyloid-related pathways in the *App^NLG-F^* model, with aging catalyzing the exacerbation of neuroinflammatory responses. The combination of these factors highlights the importance of neuroinflammation in driving disease progression in AD and suggests potential therapeutic avenues targeting these pathways.

Our study also explored how Dgat2 expression affects lipid metabolism and neurodegeneration in AD. *Dgat2*, a key enzyme involved in lipid biosynthesis, has been implicated in neuroinflammation and neurodegenerative processes (Kang & Rivest, 2012; Moraes et al., 2024). *Dgat2 KD* in Drosophila models indicate that lipid metabolism plays a critical role in locomotor, cognitive, and sleep deficits associated with AD. *Dgat2 KD* improves locomotor function, specifically in male flies expressing APP-mutant genes *(App^NLG^ and Aβ42),* where KD partially rescues climbing ability **(Figure 5a).** This suggests that *Dgat2* KD may mitigate some detrimental effects of amyloid-beta aggregation and provide protection against motor impairments seen in AD (Broussard & Brady, 2010).

Cognitive function, assessed through the olfactory aversion memory test, mirrored the locomotor results. The *Dgat2 KD* condition rescued memory impairment in both male and female AD models **(Figure 5b).** This suggests that *Dgat2* modulation can influence cognitive function, with KD providing a potential protective effect in the context of AD. Interestingly, our sensory acuity test (Meschi, Duquenoy, Otto, Dempsey, & Waddell, 2024) further confirmed these trends, reinforcing the notion that *Dgat2* manipulation, particularly KD, can help preserve cognitive function in AD models. In terms of sleep, *Dgat2* KD led to increased total sleep and reduced sleep activity at 3 weeks of age **(Figure 5c, d).** These results indicate that *Dgat2* manipulation does not drastically alter sleep behavior in younger flies, but that the effects on sleep are age dependent (Kao, Ho, Tu, Jou, & Tsai, 2020). However, in older AD flies (7 weeks), sleep patterns were largely unchanged by Dgat2 modulation, suggesting that the effects of lipid metabolism manipulation on sleep are less pronounced with age **(Figure S8, S9).** These findings highlight the complexity of lipid metabolism’s role in neurodegenerative diseases, with age and disease progression influencing the extent of *Dgat*2’s impact on sleep, cognition, and motor function.

We explored the effect of *Dgat2 KD* on lipid metabolism and neuroinflammation in *Drosophila* and *mouse* models of AD. *Dgat2* KD significantly reduces lipid accumulation, a hallmark of AD pathology, and increases synapsin levels **(Figure 6a-c).** This improvement in synapsin level supports the idea that reducing lipid accumulation may help the preservation of synaptic integrity, a crucial aspect of cognitive function in AD models (Jain et al., 2021). We also observed reduced expression of Upd1 and Upd3 cytokines associated with neuroinflammation in AD **(Figure S8)**. Interestingly, the expression of Dome and Hop receptors, which are integral to the JAK/STAT signaling pathway, remained unaffected, suggesting that *Dgat2 KD* effect on inflammation is mediated through lipid-related mechanisms, not directly via the JAK/STAT pathway (Iglesias, Morales, & Barreto, 2017).

In conclusion, both panneuronal and glial-specific expression of *App^NLG^ and Aβ42* led to severe impairments in locomotor and cognitive functions, enhanced lipid accumulation, and decreased synapsin levels. *Dgat2 KD* partially rescued these detrimental effects, providing insight into the potential therapeutic role of Dgat2 in *Drosophila.* Further research into the molecular markers involved in these processes and the role of Dgat2 modulation could provide valuable targets for mitigating neurodegeneration in the *App^NLG-F^* mouse model (**Figure 7**).

## Supporting information

none

## 4. Experimental Procedures

Due to space limitations, experimental procedures are described in the Supplementary section along with extended data in a separate PDF file.

## Author Contribution

GCM, JZ, JK, XO, and AY designed the experiments, including feedback from JJ. AY performed most sleep and all geotaxis experiments. AY performed the analysis of sleep parameters, qPCR experiments, and analysis of both *Drosophila* and mouse samples. OX performed all immunofluorescence staining of mouse brain samples. MB performed most memory assays. JCW processes all brain samples of the *Drosophila* brain and performs immunofluorescence staining and analyses with help from GCM. KM collected some sleep activity and memory data and performed immunoblot analyses. AY performed the analysis and analyzed all the data, including statistical analyses with help from GCM and JZ. AY prepared the paper with GCM’s input. All the authors reviewed the manuscript and provided their feedback.

## Acknowledgment

Stocks obtained from the Bloomington Drosophila Stock Center were used in this study.

## Data source file

All the Source data is available for this paper at source data in a single Excel sheet.

## Online Data Sharing

NA

## Abbreviations

ABCA1: ATP-Binding Cassette Transporter A1
Aβ: Amyloid β
ACC: Acetyl-CoA carboxylase
AD: Alzheimer’s disease
ApoE: Apolipoprotein E
APP: Amyloid precursor protein
BMP: Bis-phosphates
Dgat2: Diacylglycerol O-acyltransferase 2
Dgat2 KD: Diacylglycerol O-acyltransferase 2 knockdown
FAS: Fatty acid synthase
GWAS: Genome-wide association studies
PSEN1: Presenilin1
PSEN2: Presenilin2
Srebp: Sterol regulatory element binding protein
TREM2: Triggering receptor expressed on myeloid cells 2
TRF: Time-restricted feeding
WT: Wildtype

