## Supplementary material for "Regulation of lipid dysmetabolism and neuroinflammation progression linked with Alzheimer’s disease through modulation of Dgat2": none

**Supplementary Data:** A Single file contains materials and methods, ten figures and legends.

1. **Materials and Methods**

***1.1. Drosophila models***

Flies were maintained on a standard regular diet: agar 11 g/L, active dry yeast 30 g/L, yellow cornmeal 55 g/L, molasses 72 mL/L, 10% nipagen 8 mL/L, and propionic acid 6 mL/L. Flies were housed at 25 °C, 50% humidity in a 12-h light/12-h dark (LD) cycle and changed to new food every 3-4 days ^1, 2^. We used the following driver’s lines from Bloomington Drosophila Stock Center (BDSC), Elav-Gal4 (BL#458) for panneuronal expression ^3^, GLaz-Gal4 (BL#8765) for glial expression ^4^, and OK107-Gal4 (BL#854) for mushroom bodies ^5^. We used the following UAS lines: UAS-APP^NLG^ (APP human APP carrying the familial Alzheimer’s Swedish K670N, M671L, and Arctic E693G mutations, BL#33794) and UAS-Aβ42 (BL#33769). As previously described, we have also used UAS-Dgat2 RNAi (BL#107788), and UAS-GFP (BL#5431)^6-8^. As recently reported, we used the UAS-Gal4 system to drive tissue-specific expression or knockdown of Dgat2 in specific brain regions^8^. Flies with Elav> DGAT2 or Elav > Dgat2 RNAi (with a balancer) were crossed with UAS-App^NLG^, UAS-Aβ42, or control constructs to investigate their effects^47^.In addition to each UAS line driven under different drivers, the driver and UAS lines alone, and GFP overexpression serve as controls, as previously reported^6-8.^ Briefly, the progeny of the above-mentioned crossed flies will be collected at eclosion, separated by sex, held at a density of 25 flies/vial, and maintained on the diet mentioned above. As indicated in the results section, experiments were performed on 3-week-old (early middle age) and 7-week-old (old age) male or female flies as previously reported^6-8^.

***1.2. Mice***

All mouse breeding procedures and experiments were approved by the Institutional Animal Care and Use Committee (IACUC) at the University of Alabama at Birmingham. Mice were maintained in the Laboratory Animal Shared Resource with controlled air, light, and temperature conditions and fed ad libitum and free access to water. The *App^NLG-F^* mice carry three familial AD-associated mutations in the APP gene: Swedish mutation *(NL),* which enhances total *Aβ* production; Iberian mutation *(F),* which increases the *Aβ42/ Aβ40* ratio; and Arctic mutation *(G),* which promotes Aβ aggregation by facilitating oligomerization and reducing proteolytic degradation ^9^. *App^NLG-F^* and WT (controls) mice, maintained in a C57BL/6J background, were euthanized by cervical dislocation at the ages of 4 months (n=3) and 15 months (n=3). Half of the brain was used for immunohistochemistry, and half of the cortex was used for RNA analyses.

***1.3. Drosophila olfactory aversion training to test memory***

Flies were trained using the olfactory aversive conditioning method as previously described^10, 11^. Approximately 30-40 groups of flies were used for each genotype. Each genotype was exposed to two neutral odors (3-octanol and 4-methylcyclohexanol) prepared as a 1/10 dilution in mineral oil. A current of 100-V and 90-Hz was used as a shock reinforcer. Memory conditioning was performed using a T-maze apparatus (CelExplorer Labs). The olfactory aversive conditioning consists of three phases: naïve, training, and testing. During the naïve phase, flies are exposed to both odors for 3 minutes and 30 seconds to determine their odor preference based on the odor presented in each odor chamber. A single training round consists of exposing flies to odor paired with electrical shock for 2 minutes, followed by 2 minutes of exposure to odor paired without a shock. Depending on the fly’s naïve odor preference, electrical shocking will be paired with their preferred odor. After three training rounds, flies are given 10 minutes to recover. During the testing phase, flies are given 3 minutes and 30 seconds to choose between odor chambers. Once time has expired, T maze chambers are sealed, and the number of flies in each chamber is scored. A performance index (PI) was calculated based on flies’ avoidance of odor paired with electrical shock. To confirm these findings, we conducted a sensory acuity test based on avoidance index calculations, as previously described^12^.

***1.4. Drosophila locomotion assay***

To evaluate the panneuronal and glial-specific expression of APP mutation and the impact of *Dgat2 KD* on locomotor capability, we have developed a new 3D-printer-based device to analyze locomotive performance, using machine learning methods (manuscript under preparation). Briefly, the device holds 12 vials and features a Raspberry Pi Camera aimed precisely at them, which is connected to a Raspberry Pi 4B and a monitor. A custom Python script developed in our lab is running on the Raspberry Pi, controlling both the camera and the attached motor system. The video was recorded after tapping the device, and the Faster RCNN deep learning model was used to detect vial boundaries in the video. The flies' climbing behavior was recorded on video for later analysis. Followed by Python analysis processes video frames for fly detection and detection across time for average climbing were plotted.

***1.5. Drosophila sleep circadian activity***

Sleep-wake and circadian activity behavior were recorded using the Drosophila Activity Monitor (DAM, TriKinetics Inc MA, USA) system in a 12L: 12D cycle at 25°C^13^. We used 3-week and 7-week-old male and female progeny of Elav-Gal4 and GLaz-Gal4 with control lines of each of the four genes *(w^1118^, GFP, App^NLG^, Aβ42*). Drosophila activity (or wake) is measured by infrared beam crosses in the DAM system. Data was analyzed using Clock Lab and R Studio. Custom R scripts and methodology used with R Studio can be found at https://github.com/jameswalkerlab/Gill_et.al. Non-parametric One-way ANOVA with multiple comparisons, Kruskal-Wallis test for DAM system data was performed using GraphPad Prism. Drosophila sleep was defined by a period of at least 5 minutes of inactivity, demonstrated by zero beam breaks recorded. Average sleep per 24 hours (Zeitgeber Time (ZT) is a standardized way of measuring time within a circadian cycle, where ZT0 represents the beginning of the light phase and ZT12 marks the start of the dark phase), of each genotype was calculated. Five days were used for analysis of 3-week-old flies, and 3 days were used for 7-week-old fly experiments due to decreased viability in older flies. Sleep bouts were quantified by counting the number of periods of sleep as defined above. Sleep bout length was quantified by measuring the length of each sleep bout. Data for daytime sleep is from ZT0 to ZT12, and nighttime sleep is from ZT12 to ZT24.

***1.6. Histological analysis of Drosophila brain samples***

The impact of APP mutation and *Aβ42*, with and without Dgat2 modulation, on lipid and synapsin alternations was investigated as previously described^14.^ Briefly, fly heads were excised under a dissecting microscope, then fixed in 4% paraformaldehyde (PFA) in phosphate-buffered saline (PBS) for 15 minutes at room temperature with mild agitation. The heads were washed three times for 10 minutes in 1× PBS. After the final wash, heads were transferred to a 10% sucrose in PBS solution, ensuring full saturation overnight. They were next arranged in a mold using an optimal temperature (OCT) compound. Once frozen in place, the samples were sectioned at 20μm using a Leica CM3050 S cryostat and transferred onto warmed glass slides (Fisher #15-188-48). After a 30-minute drying period and application of a hydrophobic border, slides were washed three times for 5 minutes with 1× PBS to remove the dried OCT. The slides were then blocked with 3% BSA in TBS solution for 30 minutes and incubated overnight at 4°C or for 1 hour at room temperature with primary synapsin antibody (1:250, UI Developmental Studies Hybridoma Bank #3C11) in 3% BSA in TBS. After incubation, slides were washed three times for 5 minutes with PBS and incubated for 1 hour at room temperature with an AlexaFluor-750 anti-mouse fluorescent secondary antibody (1:500, Thermo Fisher # A-21037) and lipid (1:100, Lipid Spot488, Biotinum #70065). Finally, slides were washed three times for 5 minutes with 1× PBS and mounted with Antifade Mounting Medium with DAPI (0.9μg/ml, VECTASHIELD Vibrance H-1800). After overnight setting, multichannel fluorescence images were captured using an Olympus BX63 fluorescence microscope and analyzed with CellSens software at 10× magnification to view one section of the head per image.

The DAPI channel was used to define regions of interest (ROIs), as it clearly distinguishes between anatomical sub-regions in the head^14^. Multiple sections were imaged for each fly, and the average fluorescent intensity as well as average object count, and area were compiled across sections to determine the overall lipid accumulation value per group. The 488 (lipid) channel was thresholded to reduce the background signal and the minimum object size for detection was set to 20 pixels. The 750 (synapsin) channels remained thresholded to retain integrity during intensity comparison between individual subjects as well as conditions. All thresholding values and background filters were applied in batches across all images before data collection. Selection of multiple sections (n≥3) per fly and multiple flies within each condition (n≥3) ensured all areas of the head and brain are equally weighted during comparison.

***2.7. Mice brain staining, and image quantification***

One brain hemisphere was placed in 4% paraformaldehyde overnight at 4ºC, followed by sucrose incubation. The brain sample was frozen in OCT blocks and stored at -80ºC. 10μm tissue was cryosectioned. Nile red (Sigma: N3013) and LipidSpot™ 488 (Biotium NC1669425) were used for Lipid Droplet Staining following the company’s instructions. Gfap (Dako: Z0334, 1:20,000) or Iba1/AIF-1 (Cell Signaling Technology 17198S: 1:100) were paired with secondary Alexa fluor 488 (Invitrogen cat# A11008, 1:1,000). Hoechst (Fisher PI62249) was used as a nuclear stain. Images were acquired using the Keyence BZ-X810 microscope, and ImageJ was used for image analysis.

2.8. ***Real-time quantitative PCR in Drosophila and mouse brain samples***

As previously reported^8^, the heads of 3-week-old and 7-week-old flies were isolated and rapidly frozen. RNA extraction was performed using the Zymo Research Quick-RNA Microprep Kit (catalog R1051, Zymo Research, Irvine, CA, USA), which included on-column DNase I digestion. Quantitative PCR was conducted using the Sso Advanced Universal SYBR Green supermix from Bio-Rad, employing the BIO-RAD CFX Opus Real-Time PCR System. Expression levels were standardized using the 60s ribosomal protein (Rpl11) as a reference; three biological replicates were used with 8-10 flies each. Primers for qPCR are listed below:

Upd1-F: CAGCGCACGTGAAATAGCAT; Upd1-R: CGAGTCCTGAGGTAAGGGGA;

Upd2-F: AGCGTCGTGATGCCATTCA; Upd2-R: GCGATACGGATTGACATCGAA;

Upd3-F: ATCCCCTGAAGCACCTACAGA; Upd3-R: CAGTCCAGATGCGTACTGCTG;

Dome-F: CTCACGTCTCGACTGGGAAC; Dome-R: AGAATGGTGCTTGTCAGGCA;

Hop-F: CACCACCAACACCAATTC; Hop-R: GGAACGTCGTTTGGCCTTCT;

Stat92e-F: CCTCGGTATGGTCACACCC; Stat92e-R: TGCCAAACTCATTGAGGGACT;

Eiger-F: GATGGTCTGGATTCCATTGC; Eiger-R: TAGTCTGCGCCAACATCATC;

Imd-F: TCAGCGACCCAAACTACAATTC; Imd-R: TTGTCTGGACGTTACTGAGAGT;

Reaper-F: TGGCATTCTACATACCCGATCA;

Reaper-R: CCAGGAATCTCCACTGTGACT;

Hid-F: CACCGACCAAGTGCTATACG; Hid-R: GGCGGATACTGGAAGATTTGC;

Desat2-F: GTCGGCTACCCCTAGTCTGG; Desat2-F: TCGCCCTTGTGAATATGGAGT;

Srebp-F: ACCAACAGCCACCATACATCA; Srebp-R: AGACAAAGCTACTGCCCAGAG;

Dgat2-F: ATCCGTTGTGGATGGCAATG;

Dgat2-R: GGGAAAGTAATCACGATAGTGGC;

Rpl11-F: CGATCTGGGCATCAAGTACGA; Rpl11-R: TTGCGCTTCCTGTGGTTCAC;

Results are presented as 2−ΔΔCt values normalized to the expression of Rpl11 and. All reactions were performed in triplicate.

Mouse RNA extraction was performed by using RNeasy Plus Mini Kit (Qiagen, Cat#: 74134) from isolated mouse cortex samples. cDNA synthesis was performed using a High‐Capacity cDNA Reverse Transcription Kit (Thermo Fisher Scientific, Cat#: 4368814). Primers for mouse qPCR are listed below:

Bin1-F: TTCGGACCTATCTGGCTTCTG; Bin1-R: CCTCCTGAAGACACTCACTCA;

Abca7-F: AATTACACCTATCGACGGAGACA; Abca7-R: TGACGGACAGCCACTAGGA;

Epha1-F: AGGAAGTCACTCTAATGGACACA;

Epha1-R: CCTCACTCCACCCAGTCTCT;

Rhbdf2-F: GCCCACACCGTATCTGTTCTG;

Rhbdf2-R: GATGCCAGTTTTGTCGCTTGC;

Apoe-F: GACCCAGCAAATACGCCTG; Apoe-R: CATGTCTTCCACTATTGGCTCG;

Srebf1-F: TGACCCGGCTATTCCGTGA; Srebf1-R: CTGGGCTGAGCAATACAGTTC;

Eda-F: AGTGCTCAATGACTGGTCTCG; Eda-R: CGCTGCGGGGATGTAGTTTA;

Stat5b-F: CGATGCCCTTCACCAGATG; Stat5b-R: AGCTGGGTGGCCTTAATGTTC;

Ptprq-F: ATTTCTGCCACAACCTACAGC;

Ptprq-R: GGAGGGGTATTCCATGAAAGGAG;

Jak2-F: TTGTGGTATTACGCCTGTGTATC; Jak2-R: ATGCCTGGTTGACTCGTCTAT;

Ripk1-F: AGAAGAAGGGAACTATTCGCTGG;

Ripk1-R: CATCTATCTGGGTCTTTAGCACG;

Scd1-F: TTCTTGCGATACACTCTGGTGC; Scd1-R: CGGGATTGAATGTTCTTGTCGT;

Dgat2-F: TTCCTGGCATAAGGCCCTATT; Dgat2-R: CCTCCAGACATCAGGTACTCG;

Results are presented as 2−ΔΔCt values normalized to the expression of β-actin.

2.9. ***Statistical analysis***

Significance of climbing ability was determined using two-way ANOVA with multiple comparisons done with Uncorrected Fisher's LSD test. Significance was based on a chi-square test for olfactory aversion training, and the difference between fly decisions, and flies avoiding shock odor were analyzed with one-way ANOVA with Tukey’s multiple comparisons test. Lipid droplet size and density differences, and synaptic loss were performed by one-way ANOVA with Dunnett's multiple comparisons test for *Drosophila.*

Relative mean intensity of lipid, Gfap and Iba1 was measured using two-way ANOVA with Tukey’s multiple comparisons test for young and old mouse samples. For sleep activity analysis, differences between samples were determined using non-parametric one-way ANOVA with multiple comparisons done with the Kruskal-Wallis test. Quantitative PCR analyses were made using one-way ANOVA with Tukey’s multiple comparisons test performed for *Drosophila* samples, and two-way ANOVA with multiple comparisons done with Uncorrected Fisher's LSD test was performed for young/ old mouse samples. Bar graphs show mean ± SD. All statistical analyses were performed with GraphPad Prism 10. Differences were significant at values of **p* < 0.05; ***p* < 0.01; *** *p* < 0.001.

**Supplementary Figure 1**

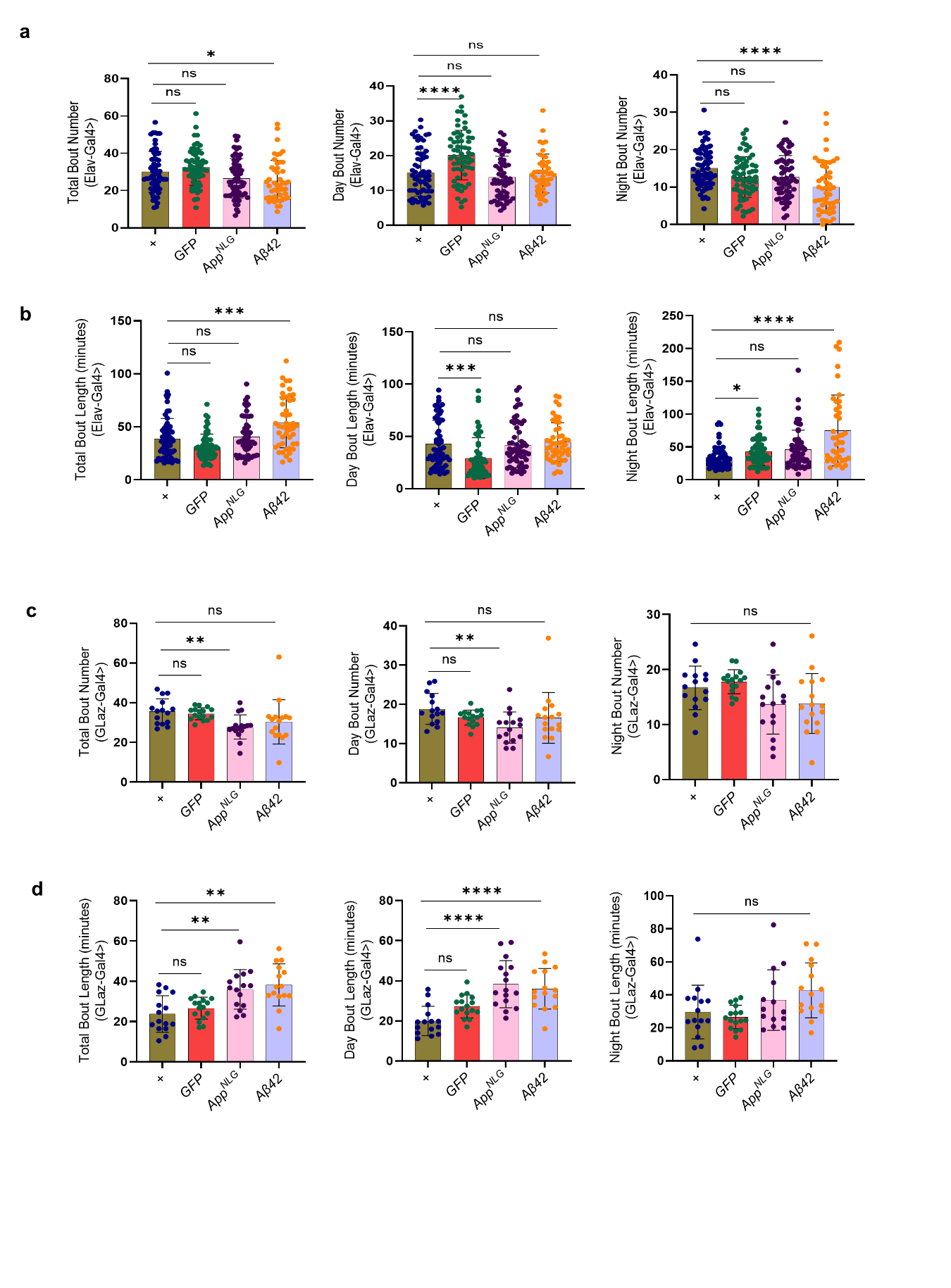

**SUPPLEMENTARY FIGURE 1. Panneuronal specific expression of *Aβ42* led to compromised sleep activity, bout number, and bout length compared to *App^NLG^* at 3-week.** (a) Total, day and night sleep bout numbers in Elav-driven *App^NLG^* and *Aβ42* models. (b) Total, day and night sleep bout lengths in Elav-driven *App^NLG^* and *Aβ42* models only compared to their respective controls. (c) Total, day and night sleep bout numbers, (d) bout length in GLaz-driven *App^NLG^* and *Aβ42* models. All experiments were performed in 3-week-old males only. Data=mean ± SD. Non-parametric One-way ANOVA with multiple comparisons, done with the Kruskal-Wallis test, was performed. Each dot represents the number of flies. “*” means *P* < 0.05; “**” means *P* < 0.01; “***” means *P* < 0.001; “ns” means not significant (an asterisk denotes significance for the average of all three replicates). Raw data and *P values* are provided in the source data.

**Supplementary Figure 2**

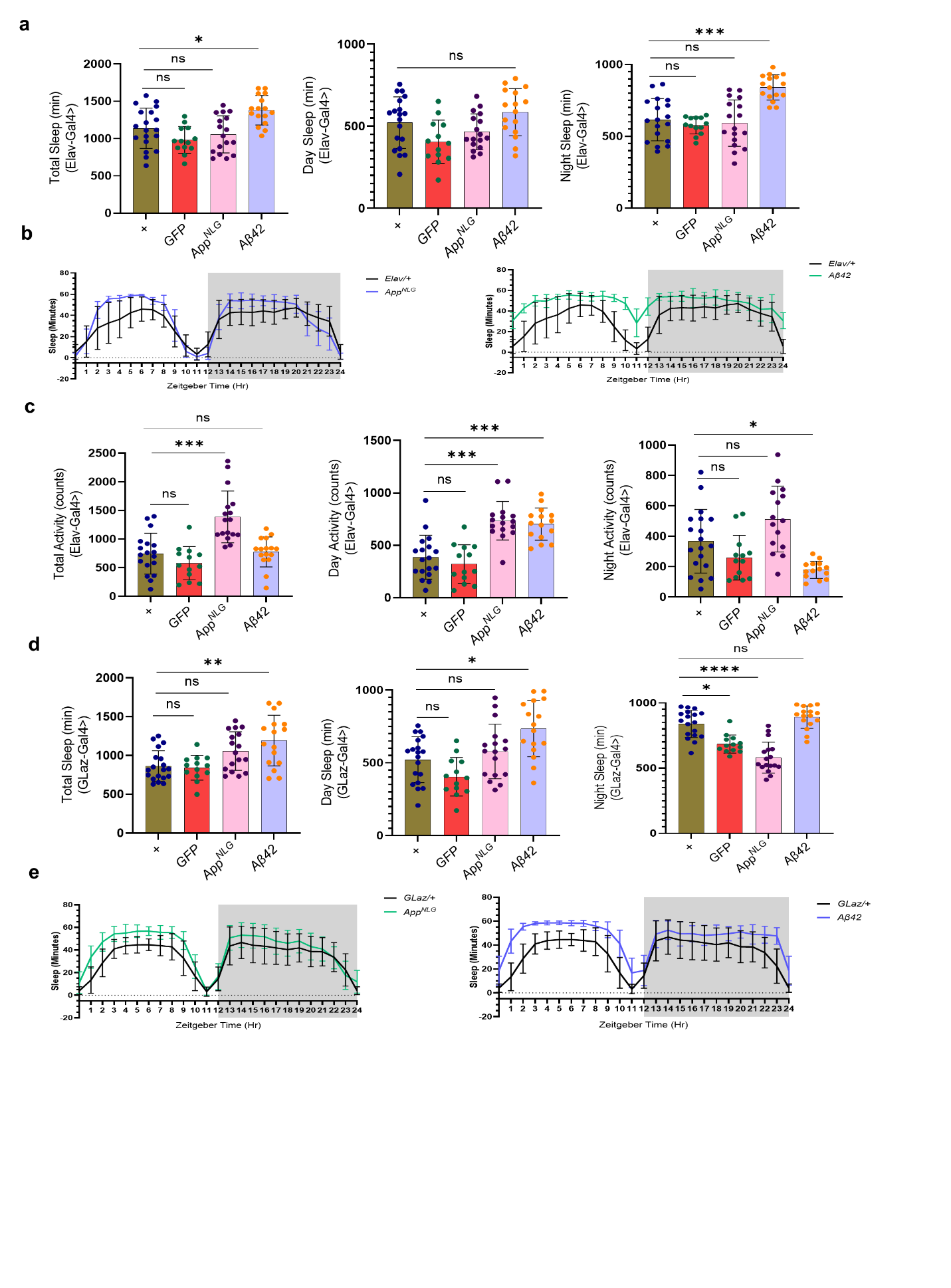

**SUPPLEMENTARY FIGURE 2.** **Panneuronal and glial-specific expression of A*β42* led to compromised seep quality compared at 7 weeks**. (a) Total, day and night sleep (in minutes) in Elav-driven *App^NLG^* and *Aβ42* models. (b) Sleep profiles at different zeitgeber time (hr.) in Elav-driven *App^NLG^* and *Aβ42* models only compared to their respective controls. (c) Total, day and night sleep activity (counts) in Elav-driven *App^NLG^* and *Aβ42* models. (d) Total, day and night sleep in (in minutes) in GLaz-driven *App^NLG^* and *Aβ42* models**.** (e) Sleep profiles at different zeitgeber time (hr.) in GLaz-driven *App^NLG^* and *Aβ42* models only compared to their respective controls. (f) Total, day and night sleep activity (counts) in GLaz-driven *App^NLG^* and *Aβ42* models. All experiments were performed in 7-week-old males only. Data=mean ± SD. Non-parametric One-way ANOVA with multiple comparisons done with Kruskal Wallis test was performed for sleep activity and sleep fragmentation. Each dot represents several flies. “*” means *P* < 0.05; “**” means *P* < 0.01; “***” means *P* < 0.001; “ns” means not significant (an asterisk denotes significance for the average of all three replicates). Raw data and *P values* are provided in the source data.

**Supplementary Figure 3**

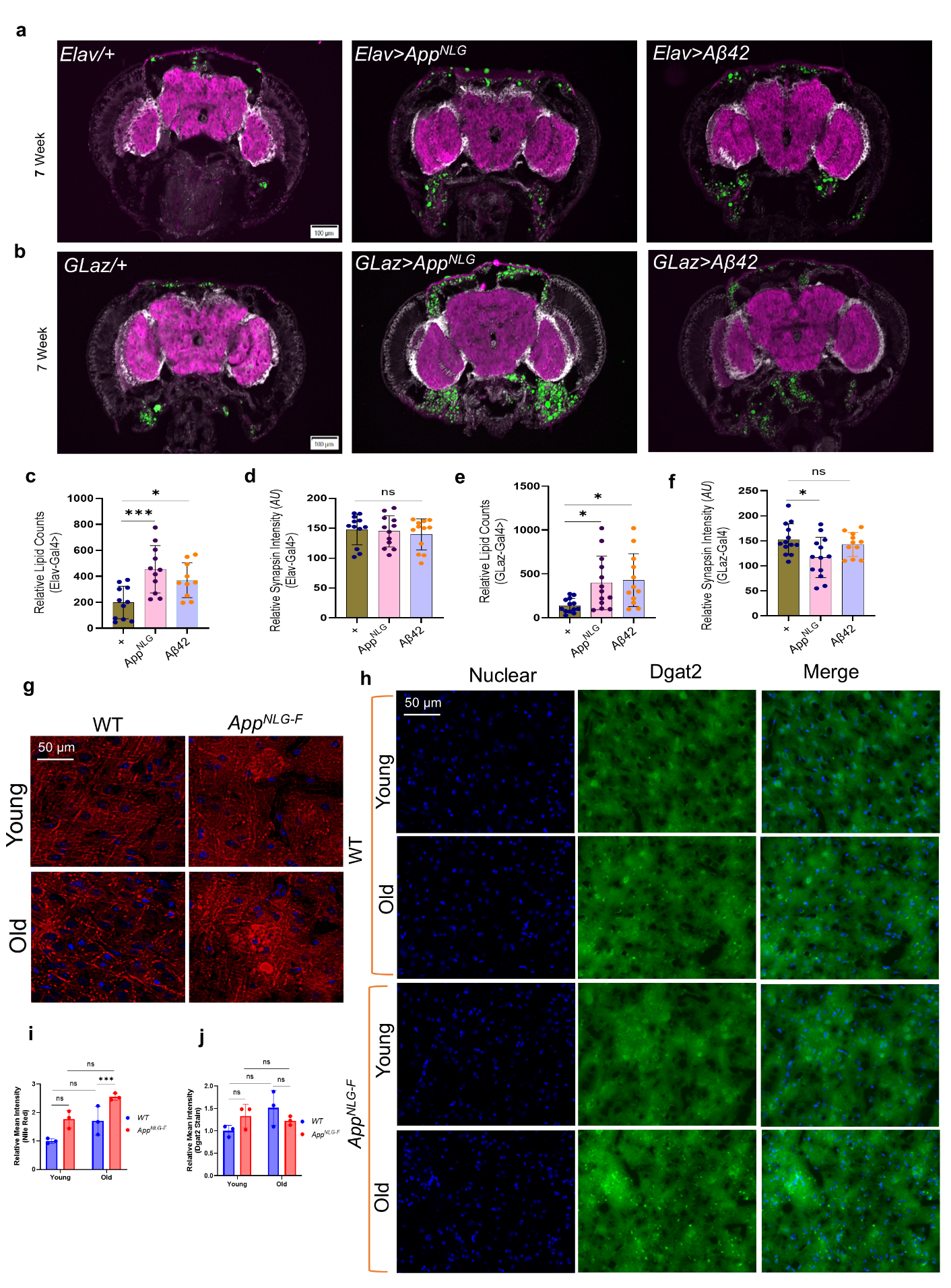

**SUPPLEMENTARY FIGURE 3. Age-dependent increase in lipid accumulation in *App^NLG^ and Aβ42* *Drosophila* models, and *App^NLG-F^* mice cortex, independent of Dgat2 expression changes.** (a, b) Representative images showing the expression of lipid accumulation (green, Lipid spot), synaptic loss (purple, anti SYNORF1), and DAPI (white), in the brains of Elav and GLaz**-**driven *App^NLG^ and Aβ42* models flies**.** (c-f) Quantification of the expression level of lipid counts and synapsin intensity in Elav and GLaz-driven *App^NLG^ and Aβ42* models. All experiments were performed in 7-week-old flies. (g) Representative image showing the intensity of Nile red (red), a marker of lipid accumulation, and DAPI (blue), in the cortical region of mouse brains from young and old wild-type (WT) and *App^NLG-F^* models. (h) Representative image showing the level of Dgat2 immunostain, and DAPI (blue), in the mouse brains of young and old wild-type and *App^NLG-F^* models**.** (i, j) Quantification of the intensity of Nile red (i) and relative mean intensity of Dgat2 (j) immunostain. Data=mean ± SD., with n=3 mice per group and 5-6 flies per group. Fold changes of fluorescence intensity were calculated relative to controls. One-way ANOVA with Tukey’s multiple comparisons test was performed for the flies’ data. Two-way ANOVA with multiple comparisons done with Uncorrected Fisher's LSD test, was performed for mouse data. “*” means *P* < 0.05; “**” means *P* < 0.01; “***” means *P* < 0.001; “ns” means not significant (an asterisk denotes significance for the average of all three replicates). Raw data and *P values* are provided in the source data.

**Supplementary Figure 4**

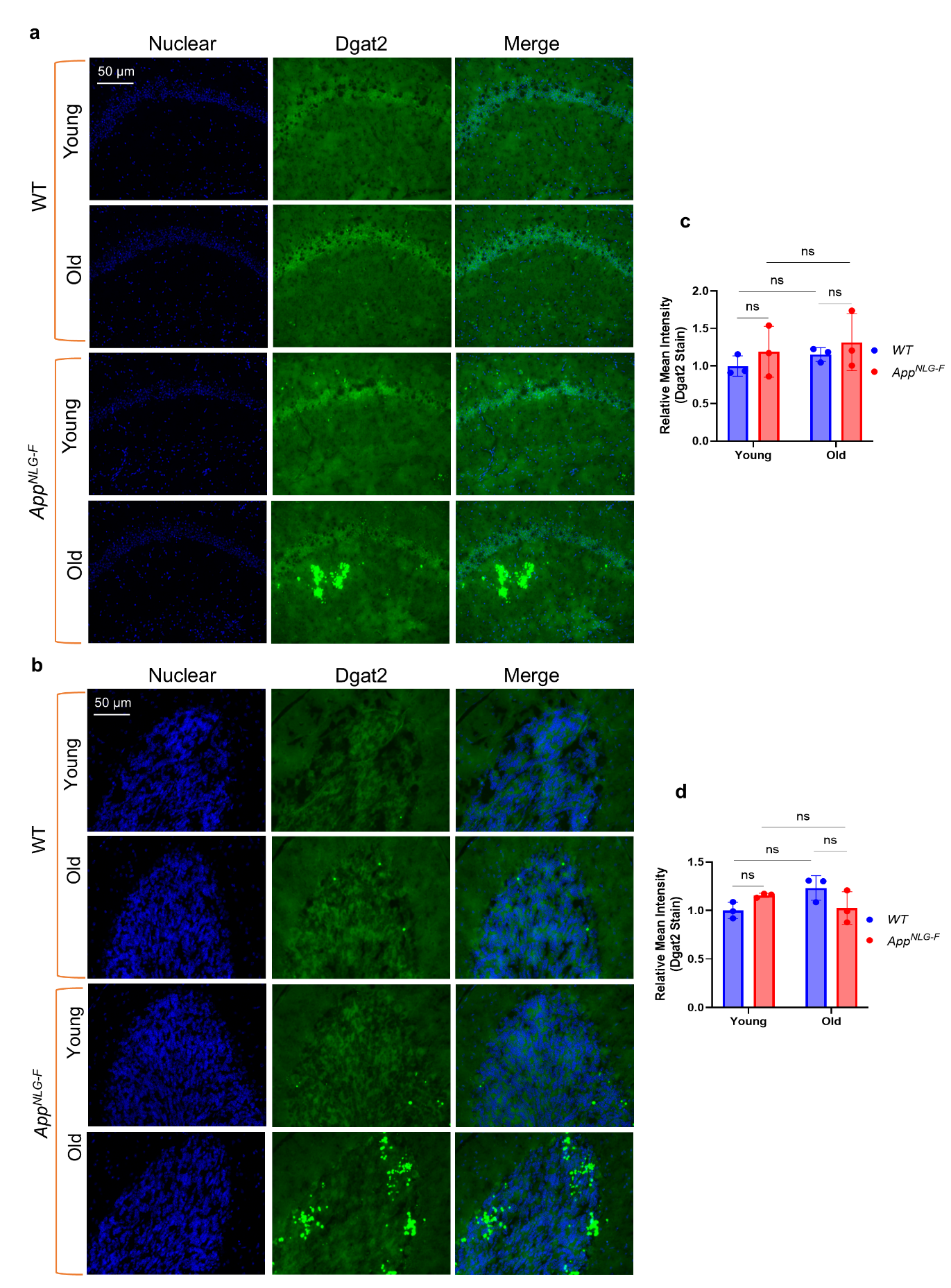

**SUPPLEMENTARY FIGURE 4.** **Unaltered Dgat2 stains in both hippocampus and cerebellum of *App^NLG-F^* mice in old age.** (a, b) Representative images showing the level of Dgat2 immunostain (green), and DAPI (blue), in the hippocampus (CA1, panel a, c) and cerebellum (CB, panel b, d) of mouse brains from young and old wild-type (WT) and *App^NLG-F^* models. (c, d) Quantification of the levels of Dgat2 immunostain in CA1 and CB**.** Data=mean ± SD., with n=3 mice per group. Fold changes of fluorescence intensity were calculated relative to controls. Two-way ANOVA with multiple comparisons done with Uncorrected Fisher's LSD test was performed for mouse data. “*” means *P* < 0.05; “**” means *P* < 0.01; “***” means *P* < 0.001; “ns” means not significant (an asterisk denotes significance for the average of all three replicates). Raw data and *P values* are provided in the source data.

**Supplementary Figure 5**

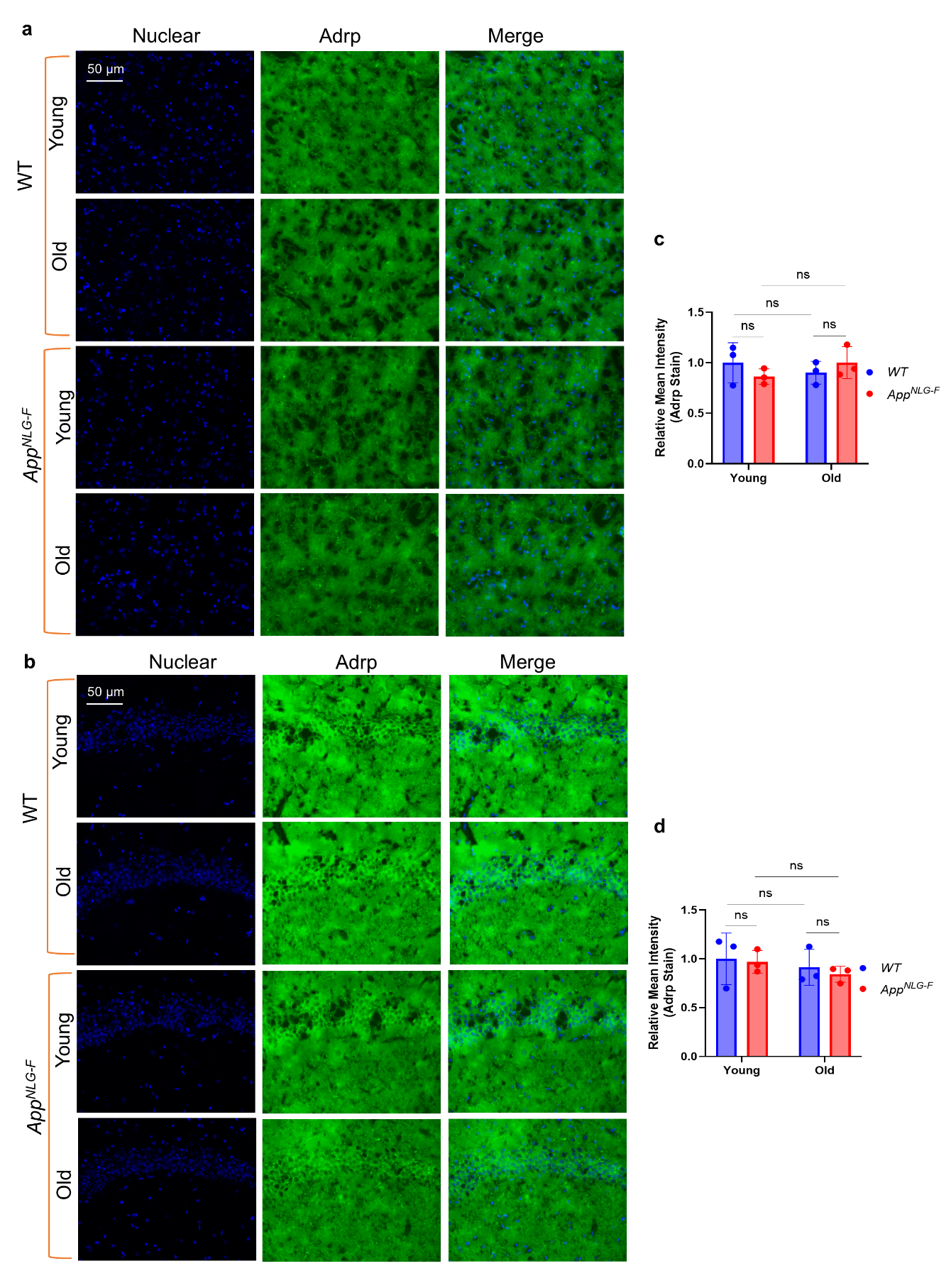

**SUPPLEMENTARY FIGURE 5. Unaltered Adrp protein in both cortex and hippocampus of *App^NLG-F^* mice in old age.** (a, b) Representative images showing the level of Adrp (Adipose differentiation-related protein) immunostain (green), and DAPI (blue), in the cortex (panel a, c) and hippocampus (CA1, panel b, d) of mouse brains from young and old wild-type (WT) and *App^NLG-F^* mice. (c, d) quantification of the level of Adrp immunostain in cortex (c) and CA1 (d). Data=mean ± SD., with n=3 mice per group. Fold changes of fluorescence intensity were calculated relative to controls. Two-way ANOVA with multiple comparisons done with Uncorrected Fisher's LSD test was performed for mouse data. “*” means *P* < 0.05; “**” means *P* < 0.01; “***” means *P* < 0.001; “ns” means not significant (an asterisk denotes significance for the average of all three replicates). Raw data and *P values* are provided in the source data.

**Supplementary Figure 6**

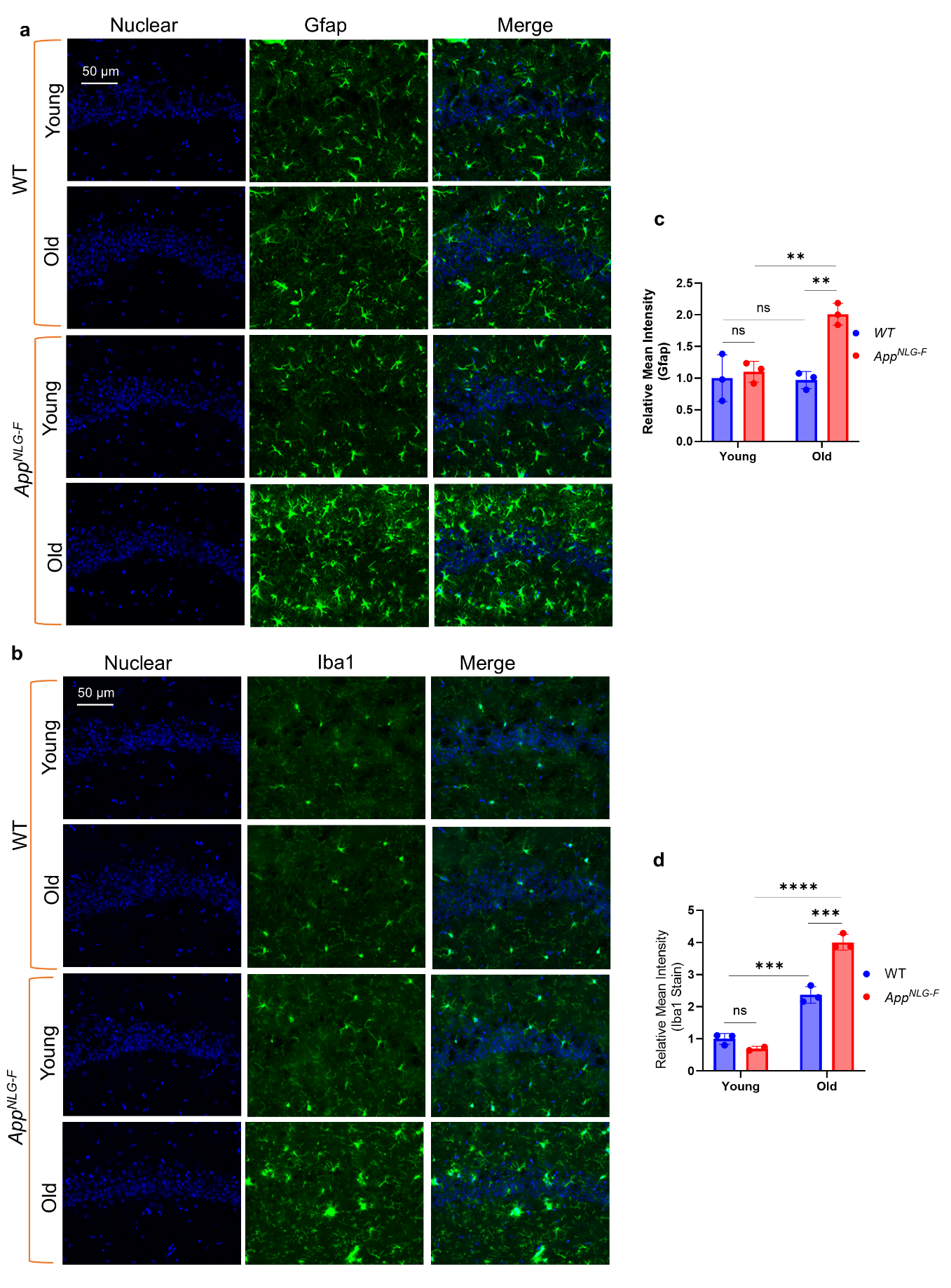

**SUPPLEMENTARY FIGURE 6. Age-dependent increase in astrocyte and microglial activation in *App^NLG-F^* mice, reflecting neuroinflammation.** (a, b) Representative images showing the levels of Gfap and Iba1 immunostain (green), and DAPI (blue), in the hippocampus of young and old wild-type (WT) and *App^NLG-F^* mice. (c, d) Quantification of the level of Gfap (c) and Iba1 (d) immunostain in the cortex. Data=mean ± SD., with n=3 mice per group. Fold changes of fluorescence intensity were calculated relative to controls. Two-way ANOVA with multiple comparisons done with Uncorrected Fisher's LSD test was performed for mouse data. “*” means *P* < 0.05; “**” means *P* < 0.01; “***” means *P* < 0.001; “ns” means not significant (an asterisk denotes significance for the average of all three replicates). Raw data and *P values* are provided in the source data.

**Supplementary Figure 7**

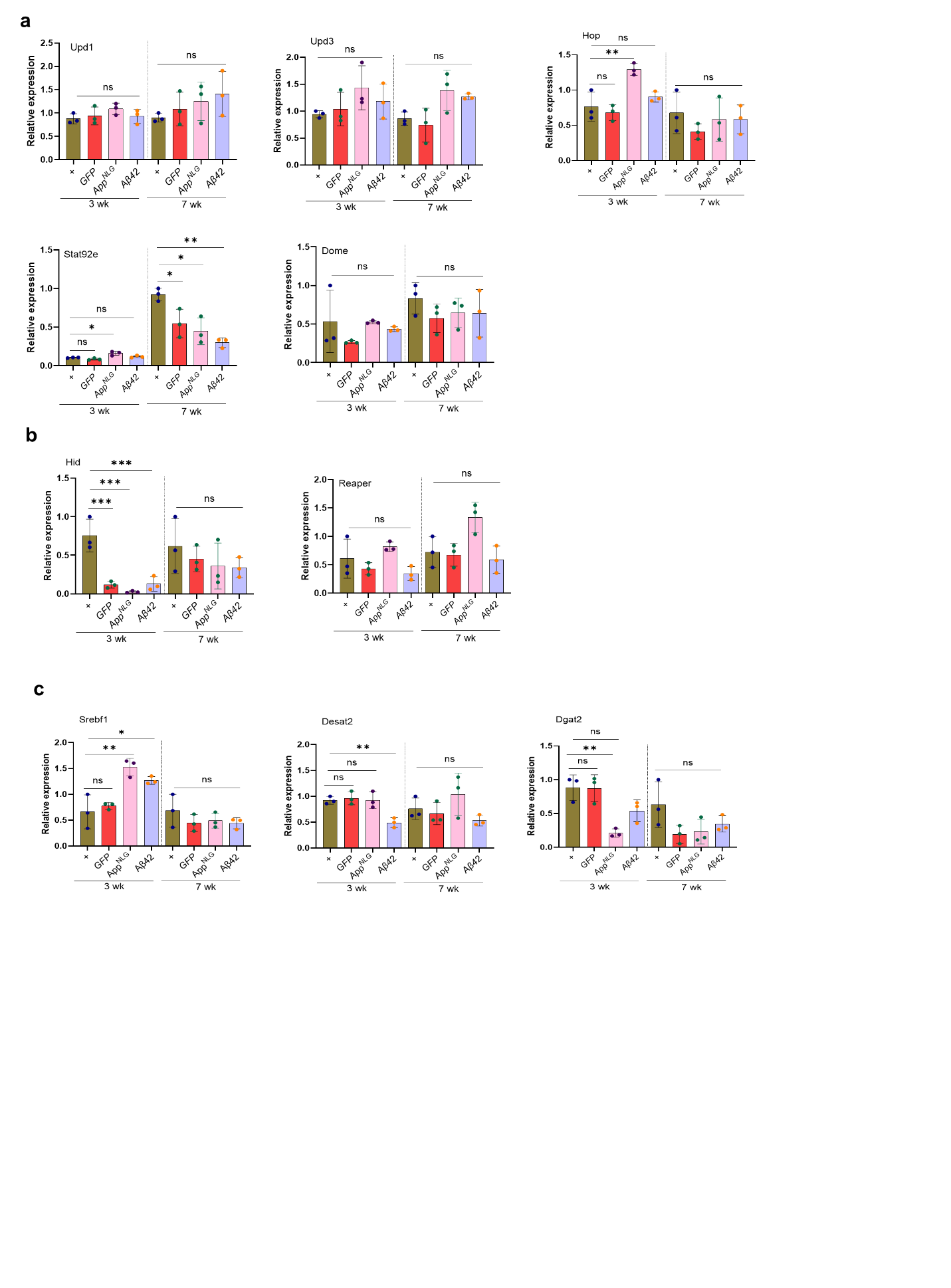

**SUPPLEMENTARY FIGURE 7. Chronic JAK/STAT signaling activation in *Drosophila* *App^NLG^* and *Aβ42* models leads to altered lipid metabolism.** (a) The qPCR result of the relative expression level of Inflammatory genes, Upd1, Upd2, Hop, Stat92e and Dome. (b) Relative expression level of cell death markers Hid and Reaper. (c) Relative expression level of Metabolic gene markers, Srebf1, Desat2 and Dgat2 in 3-week-old flies. Data=mean ± SD. One-way ANOVA with Tukey’s multiple comparisons test was performed. “*” means *P* < 0.05; “**” means *P* < 0.01; “***” means *P* < 0.001; “ns” means not significant (an asterisk denotes significance for the average of all three replicates). Raw data and *P values* are provided in the source data.

**Supplementary Figure 8**

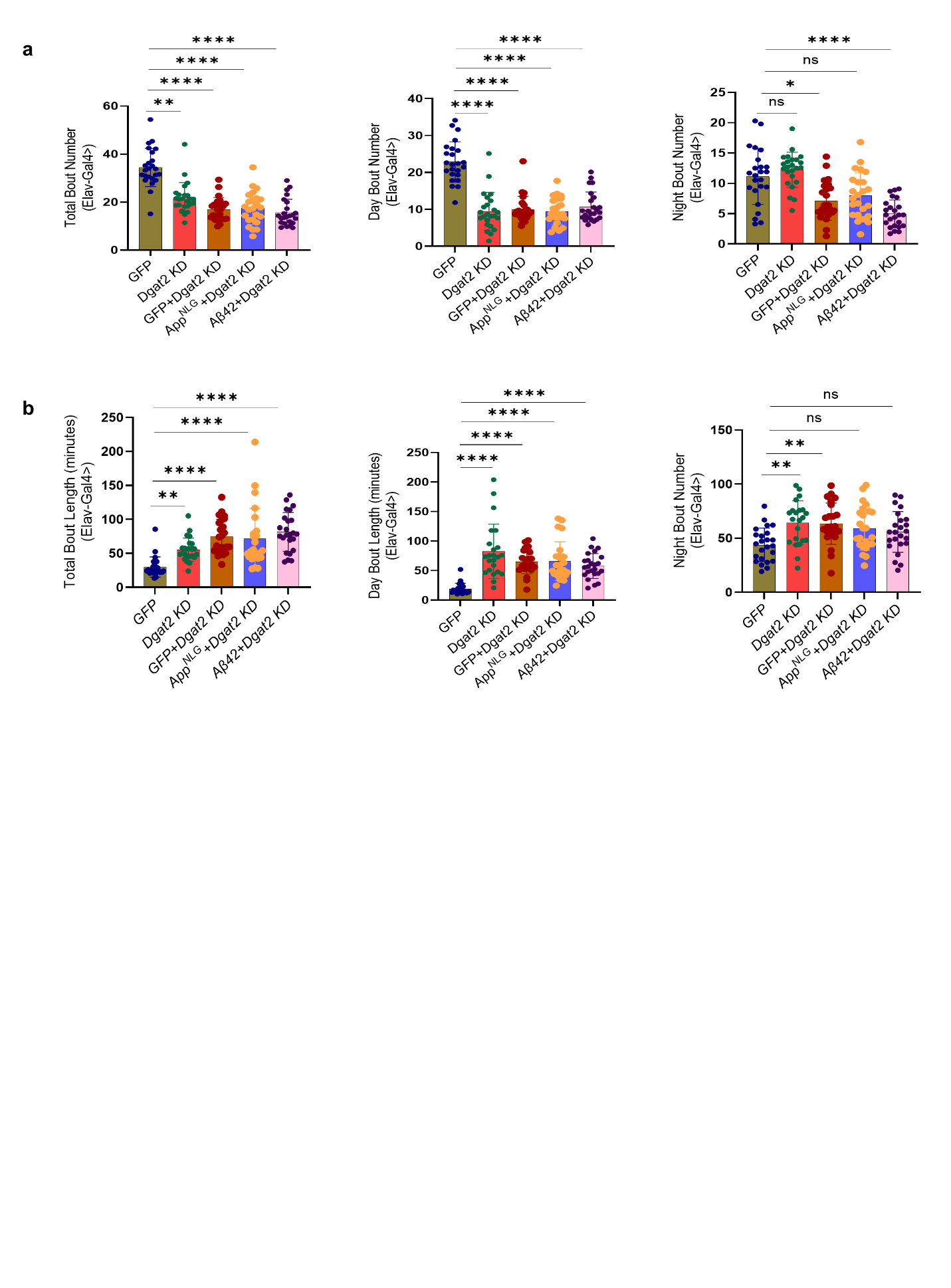

**SUPPLEMENTARY FIGURE 8. Dgat2 KD improves the sleep quality in *App^NLG^* and *Aβ42* models of flies at 3 weeks**. (a) Total, day and night sleep bout numbers. (b) Total, day and night sleep bout length in Elav-driven *Dgat KD* in *App^NLG^* and *Aβ42* models**.** All experiments were done in 3-week-old males. Data=mean ± SD. Non-parametric One-way ANOVA with multiple comparisons, done with the Kruskal-Walli’s test, was performed. Each dot represents the number of flies. “*” means *P* < 0.05; “**” means *P* < 0.01; “***” means *P* < 0.001; “ns” means not significant (an asterisk denotes significance for the average of all three replicates). Raw data and *P values* are provided in the source data.

**Supplementary Figure 9**

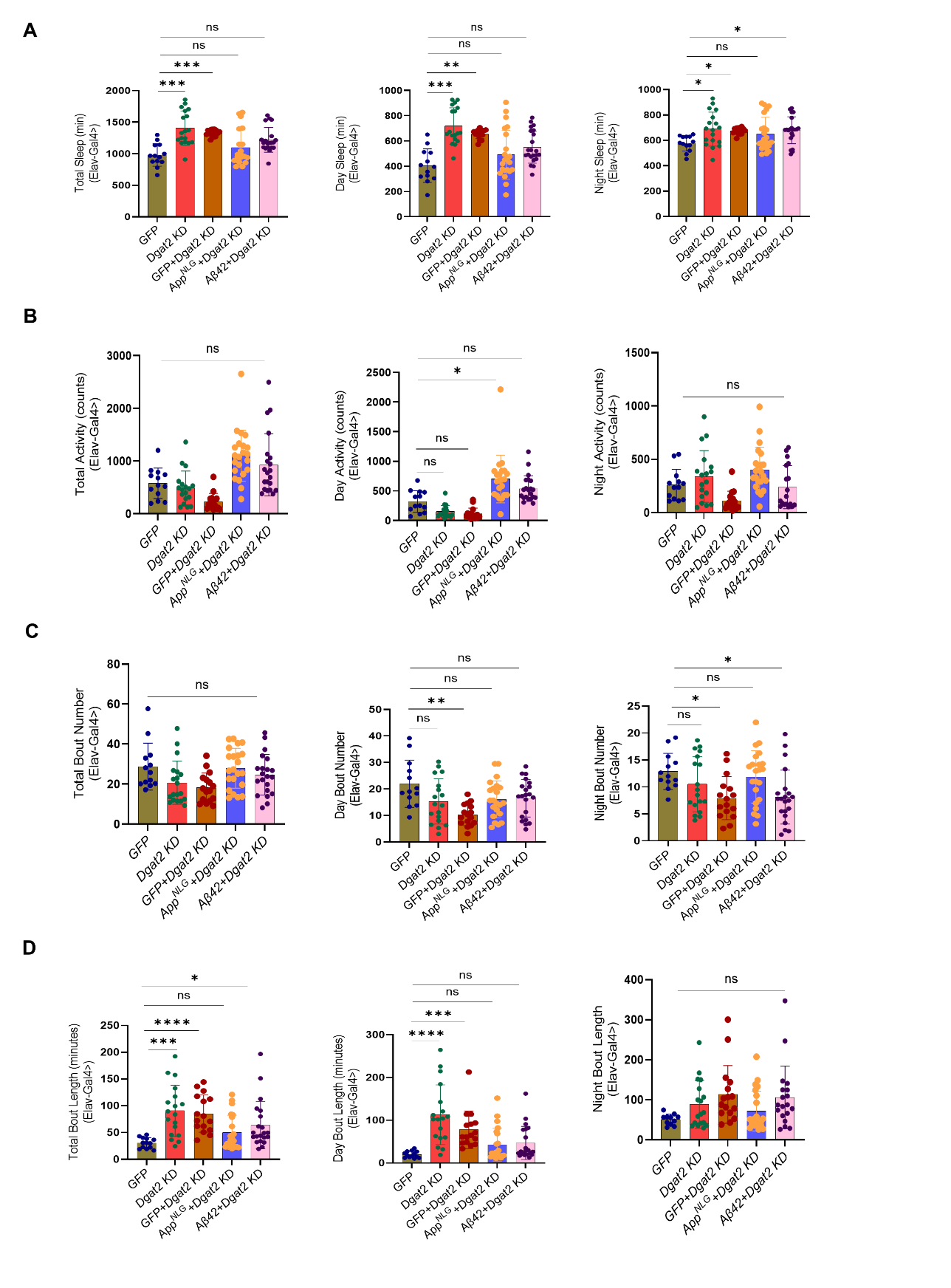

**SUPPLEMENTARY FIGURE 9**. ***Dgat2 KD* does not significantly affect sleep quality in AD models of flies at 7 weeks.** (a) Total, day and night sleep (in minutes). (b) Total, day, and night sleep activity (in counts). (c) Total, day and night sleep bout numbers. (d) Total, day and night sleep bout length in Elav-driven *Dgat KD* in *App^NLG^ and Aβ42* models at 7-week-old males**.** Data=mean ± SD. Non-parametric One-way ANOVA with multiple comparisons done with Kruskal-Wallis test was performed for Sleep parameters. Each dot represents the number of flies. “*” means *P* < 0.05; “**” means *P* < 0.01; “***” means *P* < 0.001; “ns” means not significant (an asterisk denotes significance for the average of all three replicates). Raw data and *P values* are provided in the source data.

**Supplementary Figure 10**

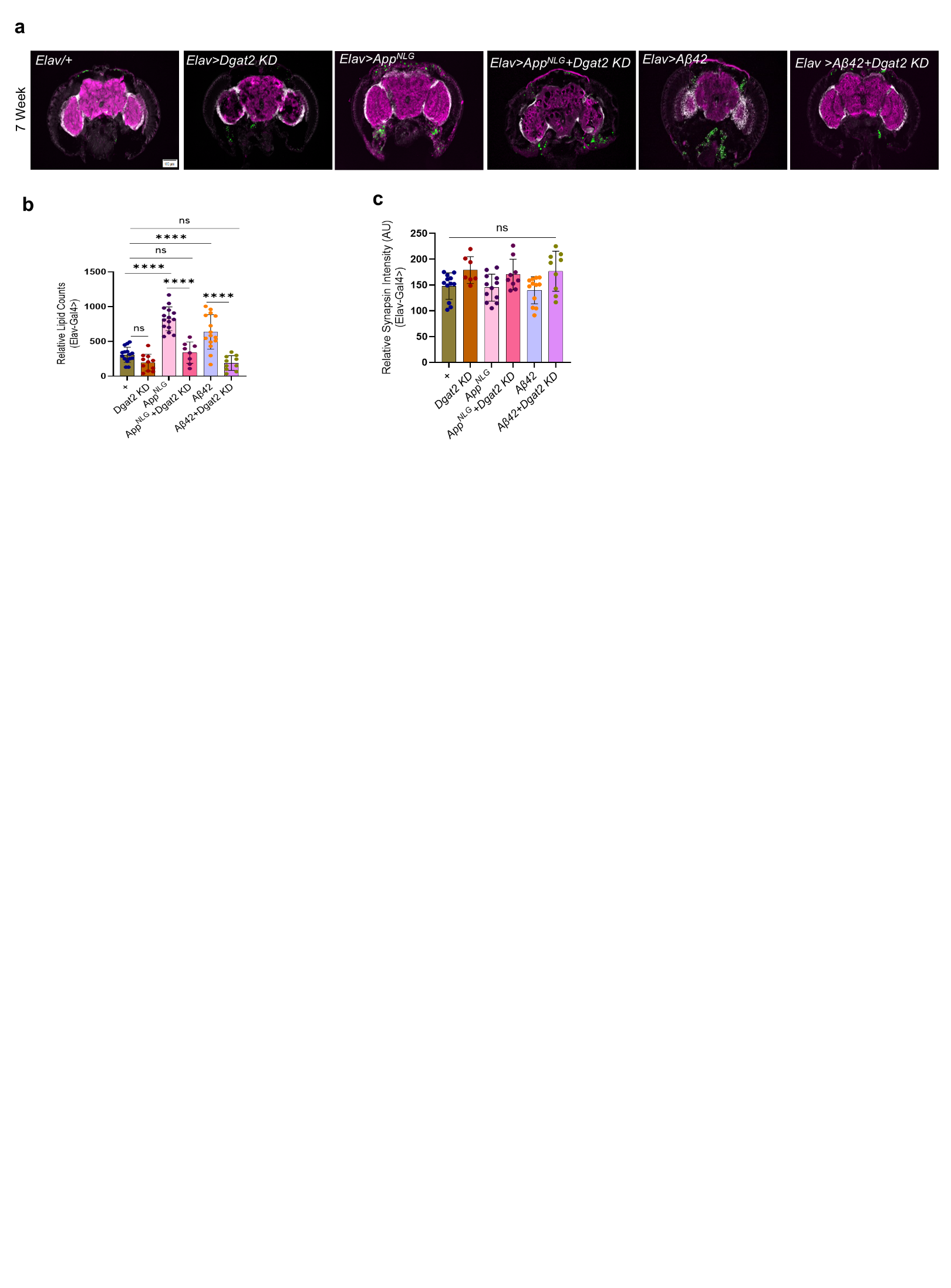

**SUPPLEMENTARY FIGURE 10.** *Dgat2 KD* reduces lipid accumulation without affecting synapsin levels in *App^NLG^ and Aβ42* *Drosophila* models. (a) Representative image showing the expression of lipid accumulation (green, lipid spot), synaptic loss (purple, anti-SYNORF1), a marker of neurodegeneration, and DAPI (white) in the brains of Elav-driven *Dgat2 KD* in *App^NLG^ and Aβ42* models. (b, c) Quantification of the expression level of lipid counts and synapsin intensity in Elav-driven *Dgat2 KD* in *App^NLG^ and Aβ42* models. All experiments were performed in 7-week-old flies. Data=mean ± SD., with n=5-6 flies per group. Fold changes of fluorescence intensity were calculated relative to controls. One-way ANOVA with Tukey’s multiple comparisons test was performed for flies. “*” means *P* < 0.05; “**” means *P* < 0.01; “***” means *P* < 0.001; “ns” means not significant (an asterisk denotes significance for the average of all three replicates). Raw data and *P values* are provided in the source data.
